# Alanine tRNA translate environment into behavior in *Caenorhabditis elegans*

**DOI:** 10.1101/289629

**Authors:** Diana Andrea Fernandes De Abreu, Thalia Salinas-Giegé, Laurence Drouard, Jean-Jacques Remy

**Affiliations:** Genes, Environment, Plasticity, UMR CNRS 7254, INRA 1355, Université Nice Côte d’Azur, 06903 Sophia-Antipolis, France.; Institut de biologie moléculaire des plantes-CNRS, Université de Strasbourg, F-67084 Strasbourg, France

## Abstract

*Caenorhabditis elegans* nematodes produce and maintain imprints of attractive chemosensory cues to which they are exposed early in life. Early odor-exposure increases adult chemo-attraction to the same cues. Imprinting is transiently or stably inherited, depending on the number of exposed generations.

We show here that the Alanine tRNA (UGC) plays a central role in regulating *C. elegans* chemo-attraction. Naive worms fed on tRNA^Ala^ (UGC) purified from odor-experienced worms, acquire odor-specific imprints.

Chemo-attractive responses require the tRNA-modifying Elongator complex sub-units 1 (*elpc-1*) and 3 (*elpc-3*) genes. *elpc-3* deletions impair chemo-attraction, which is fully restored by wild-type tRNA^Ala^ (UGC) feeding. A stably inherited decrease of odor-specific responses ensues from early odor-exposition of *elpc-1* deletion mutants.

tRNA^Ala^ (UGC) may adopt various chemical forms to mediate the cross-talk between innately-programmed and environment-directed chemo-attractive behavior.

## INTRODUCTION

Although parental adaptation to different environmental challenges can be transmitted to future generations, the mechanisms by which external signals are translated into heritable information are largely unknown (***Liberman et al., 2019***). *C. elegans* worms keep a life-term memory of attractive olfactory cues to which they were exposed during the first larval stage (***Remy and Hobert, 2005***). Early odor-exposure results in a significant enhancement of odor-specific chemo-attraction at the adult stage. Such olfactory imprinting can be inherited either transiently by a single generation or stably over generations after at least five consecutive generations were exposed to the same cue (***Remy, 2010***).

Non-coding RNAs have previously been implicated in the memory and transgenerational transmission of different environmental informations both in the *C. elegans* nematode (***Rechavi et al., 2014***; Juang et al., 2013; Hall et al., 2013; ***Posner et al., 2019***) and in the mouse (***Gapp et al., 2014***; Grandjean et al., 2015; ***Benito et al., 2018***).

In this work, we found that a single Alanine tRNA molecule plays an essential role in regulating both the innately expressed and the environment modulated chemo-attractive responses in *C. elegans*. We first observed that naive unexposed worms acquire odor-specific imprinting after they were fed on RNA extracted from odor-exposed animals. Biochemical fractionation of RNAs led to the identification of the transfer RNAs (tRNAs) containing fraction as the imprinting transmission medium. Using tRNA-specific probes, we found that among all *C. elegans* tRNA molecules, only the Alanine tRNA with the UGC anti-codon (tRNA^Ala^ (UGC)), transfers odor-specific imprints to naive worms via feeding. The highly purified tRNA^Ala^ (UGC) from worms exposed to three different attractive odors - benzaldehyde (BA), citronellol (CI) and isoamyl alcohol (IA) - transfers the same odor-specific heritable behavioral changes as does early odor stimulations. This single tRNA molecule could thus carry different odor-specific codes, according to the odors worms were exposed.

The nucleotides of all forms of coding and non-coding RNAs can be chemically modified. To date, more than 160 different chemical modifications of RNA bases have been described. Each modification involves specific reactions catalysed by enzymes called « writers » (Sarin and Leidel, 2014; Schaefer et al., 2017; Jonkhout et al., 2017; ***Boccaletto P. et al., 2018*).**

It is largely admitted that, compared to all other forms of RNAs, transfer RNAs are the most extensively modified. The combination of modified bases would potentially produce a great number of chemical variants for a single tRNA molecule.

We hypothesized that specific patterns of base modification makes the odor-codes written on tRNA^Ala^ (UGC) molecules upon odor-stimulation. We therefore analysed the behavioral effects of mutations inactivating the Elongator complex, ELPC, the only known tRNA bases modifyer known in the nematode. Inactivation of the elongator complex sub-units 3 (*elpc-3*) or 1 (*elpc-1*) differently impairs the chemo-attractive behavior.

*elpc-3* deleted worms are not anymore attracted by CI, BA or IA. Chemo-attraction is however fully rescued to wild-type levels after feeding *elpc-3* mutants on naive wild-type tRNA^Ala^ (UGC). This suggests *elpc-3* is required for the synthesis of the chemical form of tRNA^Ala^ (UGC) which support the development of chemo-attractive responses.

Furthermore, the absence of a functional ELPC-1 in the *elpc-1* deleted mutants reverses the outcome of odor-exposure: as opposed to wild-type, exposure stably decrease adult odor-specific responses in *elpc-1* mutants and their progeny. Such negative odor-specific imprinting is impaired upon addition of odor-specific amounts of a chemically modified Uridine in worm food. Epitranscriptomic regulation of tRNA^Ala^ (UGC) appear as essential molecular links between environmental inputs and stably inherited changes of the *C. elegans* chemo-attractive behavior.

## RESULTS

Olfactory imprinting is a long-term inherited behavioral change induced by the early olfactory environment. We hypothsized RNAs could convey the olfactory imprints. First, olfactory imprinting was induced, as previously described (***Remy and Hobert, 2005***), by exposing animals to two different olfactory cues, citronellol (CI) or benzaldehyde (BA) (*Figure 1a*, odor-exposure). The efficiency of this induction was tested using the chemotaxis assay described in Methods and in ***Supplement Figure Method 1***. second, total RNA was extracted from CI-exposed, BA-exposed or water-exposed control worms. Third, these RNAs were fed to naive larvae (***Figure 1a***, RNA-feeding). Fourth, once RNA-fed larvae reached the adulthood, they were subjected to the chemotaxis assays. We observed the naive worms fed on RNA from CI or BA-exposed worms migrate significantly faster toward CI or BA, as if they had been themselves odor-exposed (***Figure 1b***).

**Figure 1.**
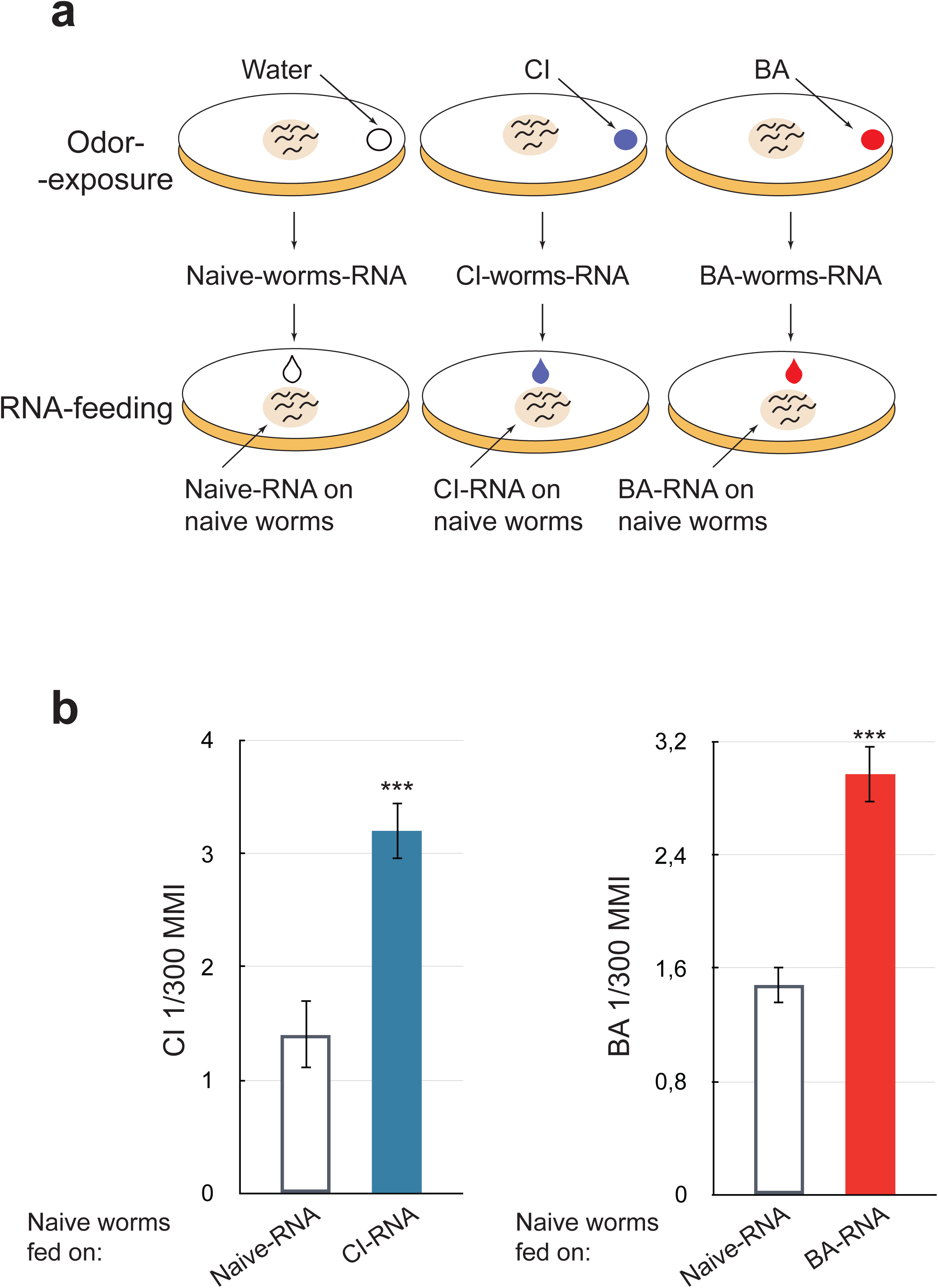
RNA extracted from odor-exposed worms transfer olfactory imprinting to naive unexposed worms via feeding. (1a) Schematic depiction of worms odor-exposure and RNA-feeding. Upper part: a drop of 4l of citronellol (CI) or benzaldehyde (BA) or water (for control⎧unexposed) is placed on the lid of each culture plate. Worms produce long-lasting odor-specific imprints when exposed to odors for 12 hours post-hatch at 20°C, the critical period for olfactory imprinting (*Remy and Hobert, 2005*). Worms exposed to odors during this period are collected at the adult stage. RNA is extracted from each collected population. Bottom part: a 10l drop of purified RNA is placed on worm⎧ food (*E. coli* 0P50) to be ingested by naive larvae. **(1b)** Chemotaxis assays performed on naive adults fed on different RNA populations. Worms fed on RNA from CI-exposed (CI-RNA, blue column) and worms fed on RNA from BA-exposed (BA-RNA, red column) migrate faster toward a CI source and a BA source, respectively, compared to worms fed on RNA from naive animals (Naive-RNA, white columns). Mean Migration Index (MMI) was determined as described in Supplemental Material (experimental repeats > 4, ***p-value<0.001).

We observed that dor-specific imprinting can be elicited by the exogenous addition of RNA extracted from odor-exposed worms to the food of naive unexposed worms. This observation shows that worms exposed to odors during the L1 larval stage produce RNA populations able to alter chemosensory responses of naive animals via ingestion.

To identify which RNA molecules are able to transfer odor imprints, we started by separating the large from the small RNAs. We studied the transfer of imprinting by feeding naive worms on either the “large RNAs fraction” or the “small RNAs fraction”. We observed that only RNAs smaller than 200 nucleotides (nt) are able to trigger the imprinting (Supplemental ***Figure 1a***). Migration on a 3.5% agarose gel separates small RNAs into five fractions (A to E bands, insert on Supplemental ***Figure 1b***). After RNA-feeding, we observed that olfactory imprints were exclusively transmitted by the “D” small RNA fraction. Based on co-migration with a double-stranded RNA ladder (L), the “D” RNA population migrates with an apparent mean size of 45 nt on this non-denaturing electrophoresis condition.

Further fractionation of the imprinting RNA fraction on denaturing 7 M urea-15 % polyacrylamide gel (***Figure 2a***) revealed it is composed of several RNA species we named fraction 1 to fraction 8. We noticed that fractions 2 to 7 represent a typical profile of transfer RNAs (tRNAs) (***Figure 2a, left panel***). Performing northern-blot analyses using two *C. elegans* specific probes for tRNA^Leu^ (AGG) and tRNA^Gly^ (UCC), we confirmed that fractions 2 to 7 indeed contain the *C. elegans* tRNA population (***Figure 2a, right panel***).

**Figure 2.**
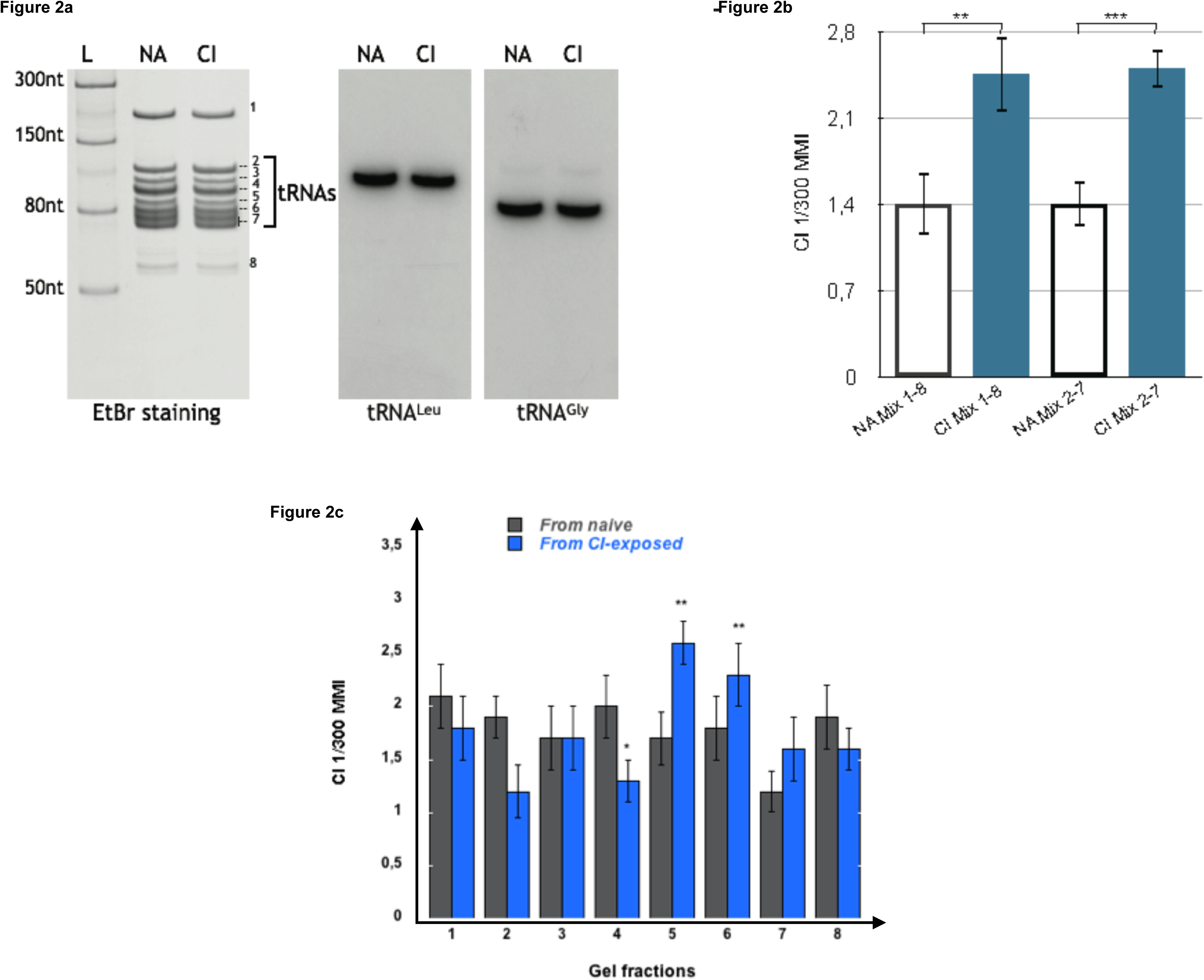
Olfactory imprinting is triggered by the transfer RNAs (tRNAs) containing fraction. **(2a)** Left part: Total small RNA from naive (NA) and from CI-exposed (CI) were fractionated on a 7M Urea 15% polyacrylamide gel and stained with ethidium bromide (EtBr). Band 1 is the 5S rRNA, bands 2 to 7 are tRNAs, while band 8 is made of unknown RNAs. The L well contains the NEB single stranded RNA ladder. Right part: Northern blot analyses were performed on NA and CI RNA blots with two radiolabeled probes specific to the *C. elegans* tRNA^Leu^(AAG) and tRNA^Gly^(UCC). **(2b)** After elution from the polyacrylamide gels, fractions 1 to 8 or fractions 2 to 7 from naive (NA Mix 1-8 and NA Mix 2-7) and from CI-exposed (CI Mix 1-8 and CI Mix 2-7) were pooled. Naive worms were fed on each pool and assayed for CI response. Both of the pooled CI mixes trigger a faster migration (MMI) in CI gradients than the corresponding NA mixes (experimental repeats > 4, **p=0.006; ***p<0.001). **(2c)** Naive N2 were fed on each of the 1 to 8 individual fractions from naive and from CI-exposed. Only feeding on the tRNA fractions 5 and 6 from CI-exposed enhances migration in CI gradients (MMI), compared to all other tRNA fractions (experimental repeats > 4, ***p<0.001).

We next wanted to know if the imprinting activity is spread over the whole tRNA co-purified population or linked to specific fractions. We cut and eluted each of the indicated 1 to 8 fractions from the polyacrylamide gels. We reconstituted the whole imprinting fraction by mixing fractions 1 to 8, and the whole tRNA co-migrating RNA populations by mixing fractions 2 to 7. Both 1 to 8 (CI Mix 1-8) and 2 to 7 (CI Mix 2-7) mixes extracted from CI-exposed worms, transferred a CI-specific imprint to naive worms, while the corresponding NA mixes (NA Mix 1-8 and NA Mix 2-7) did not (***Figure 2b***).

After a detailed fraction by fraction analyses, we found that feeding naive on fractions 5 and 6 significantly enhances CI chemo-attraction (***Figure 2c***).

596 functional tRNA genes have been predicted in the *C. elegans* genome (***Duret, 2000; Robertson and Thomas, 2006; Chan and Lowe, 2009***). To be fully active, tRNAs need to be heavily modified post-transcriptionally (***Agris, 2015; Duechler et al., 2016***). Indeed, the high abundance of tRNA post-transcriptional modifications blocks the progression of reverse-transcriptases, thus cDNA synthesis, making quantitative tRNA sequencing very challenging. In order to obviate these limitations, alternative methods have been proposed in the recent literature (***Pang et al., 2014; Zheng et al., 2015; Cozen et al., 2015; Shigematsu et al., 2017***), however none of them is readily available. For this reason, instead of a sequencing strategy, we used biochemical methods to identify which RNAs molecules are able to transfer olfactory imprinting among these tRNA co-purified fractions.

To assess if some tRNAs are responsible for imprinting, we combined streptavidin microbeads purification with northern blot analysis. According to the genomic tRNA database (GtRNAdb) predictions, *C. elegans* uses 46 different anticodons to decode the 20 aminoacids. Five anticodons are used for Ser, Arg and Leu, three for Ala, Gly, Pro, Thr and Val, two for Lys, Glu, Gln and Ile, and one for Phe, Asp, His, Met, Tyr, Cys and Trp.

As mentioned above, imprinting is transferred only by fractions 5 and 6. tRNAs bearing different anticodons decoding the same amino-acid, named tRNA isoacceptors, display a high degree of sequence homology. Since the denaturing gel shown in ***Figure 2a*** separates tRNA molecules on the basis not only of their length but also of their nucleotide sequences, we hypothesized that each fraction, including the imprinting fractions 5 and 6, may contain a limited number of isoacceptor tRNAs. For microbeads purifications, we used a set of 37 nt long oligonucleotides, specific to each tRNA isoacceptors family and complementary to their respective tRNAs 3’ halves, as described in Methods. Except two out of the five decoding Ser and Arg and one out of the five decoding Leu, all anticodon-specific tRNAs used by *C. elegans* are represented in our set of tDNA probes. For northern blots analysis, we used short aminoacid-specific isodecoder tRNAs probes (Methods).

We reasoned that if a mixture of tRNAs eluted from a pool of tDNA oligonucleotides bound to microbeads transfers imprinting, then this population should necessarily contain the imprinting RNAs.

tRNAs were purified from odor-exposed (CI or BA) worms using 14 different pools made of oligonucleotides specific to 7 to 11 different tRNA ***(A to N, Table 1)***. The pools were designed such as each isotype (amino-acid decoding) tRNA is present in three different pools. Out of the 14 pools, only the A, J and N pools purified tRNAs were able to transfer the imprinting.

Pool A is made of oligonucleotides specific to Ala, Arg, Asn, Asp and Cys tRNAs. Arg, Asn, Asp and Cys oligonucleotides did not allow the purification of the imprinting tRNAs, as they are also part of the imprinting negative pools F and K. Pool J is made of the Tyr, Val, Ala and Arg probes, but Tyr and Val probes are part of the imprinting negative pool E.

Pool N is made of the Thr, Trp, Tyr, Val and Ala probes, but Thr, Trp, Tyr and Val probes are part of the imprinting negative pools E and I.

Altogether, we concluded that none but the Ala oligonucleotides present in the three imprinting positive pools A, J and N purified the imprinting tRNAs.

We further proceeded with tRNA purification by isotype-specific probes, and confirmed that only the tRNAs eluted from Alanine tDNA probes imprint naive worms, as shown in ***Figure 3a***. It remains possible that other RNA species that co-elute from the Alanine tDNA probe are the active species.

**Figure 3 and Table 1.**
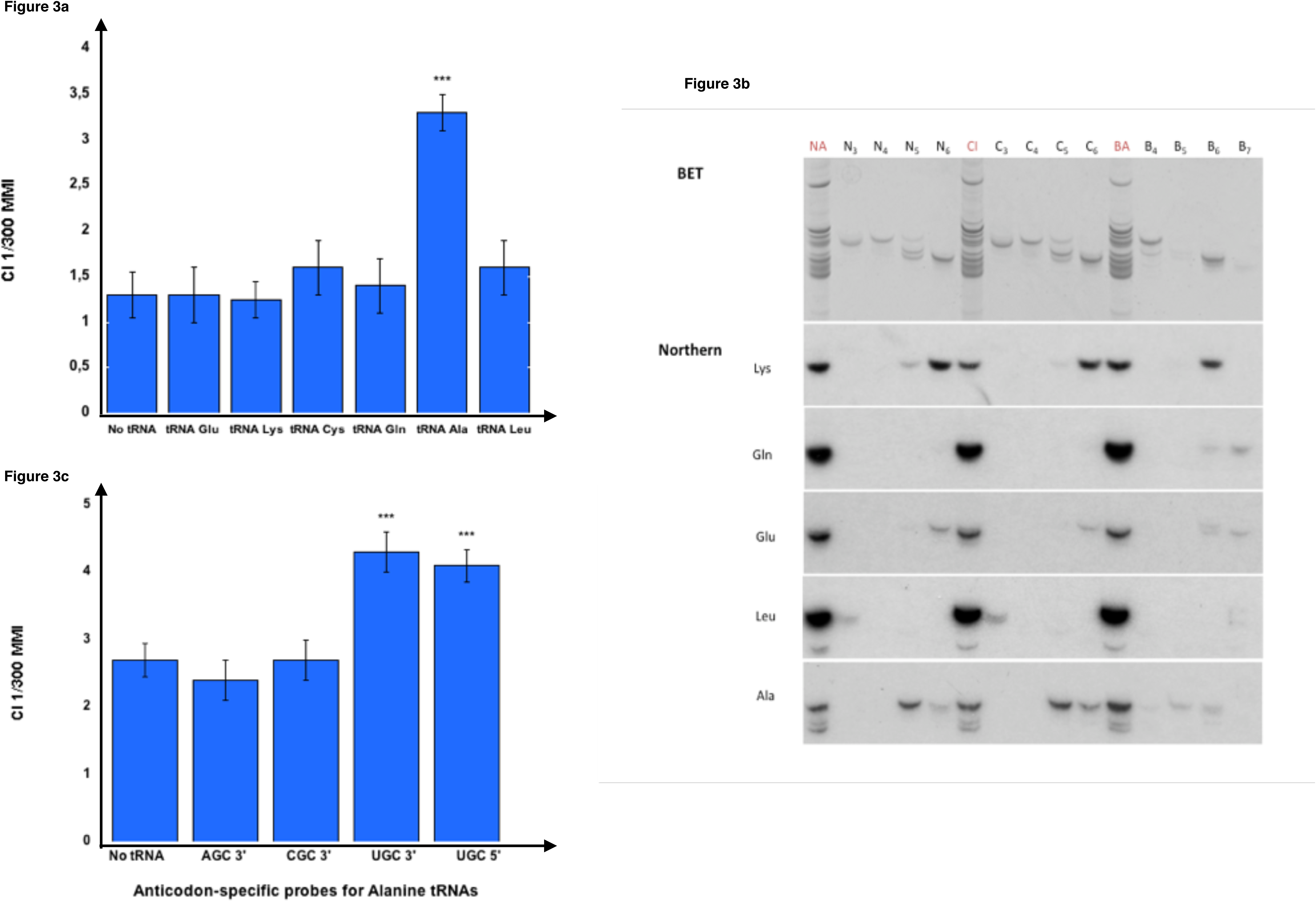
Olfactory imprinting is transferred by Alanine tRNA (UGC) **(3a)** 37 nt long isotype and isoacceptor-specific DNA probes corresponding to the 3’ halves of *C. elegans* tRNAs were designed (Methods). Probes were 3’-biotinylated and used for purification on streptavidin microbeads, as described in the Methods. Small RNAs or gel-eluted fractions 5 or 6 from odor-exposed worms were hybridized to pools of isotype-specific probes bound to microbeads. The tRNA eluted from each pool was assayed to test the imprinting transfer to naive. The CI imprinting Fraction 5 CI-tRNAs were hybridized to the isotype-specific 3’-biotinylated DNA probes for the 3’ halves of tRNA^Glu^, tRNA^Lys^, tRNA^Cys^, tRNA^Gln^, tRNA^Ala^ or tRNA^Leu^ bound on streptavidine-microbeads. Naive worms were fed on tRNAs eluted from the microbeads and tested for migration in CI 1/300 gradients (***p-value < 0.001). **(3b)** Northern blot analysis on gel-eluted tRNA fractions from Naive (N3 to N6), on tRNA fractions from CI-exposed (C3 to C6) and on tRNA fractions from BA-exposed (B4 to B7) N2 worms. Hybridizations were performed with radiolabeled oligonucleotides specific for tRNA^Lys^ (UUU), tRNA^Gln^ (UUG), tRNA^Glu^ (UUC), tRNA^Leu^ (AAG), and tRNA^Ala^ (AGC, CGC, UGC). **(3c)** Fraction 5 CI-tRNAs were eluted from microbeads bound to the four Alanine codon-specific (3’-AGC, 3’-CGC, 3’-UGC and 5’-UGC) probes. Naive worms were fed on tRNAs eluted from the microbeads and tested for migration in CI 1/300 gradients (***p-value < 0.001).

**Table 1.**
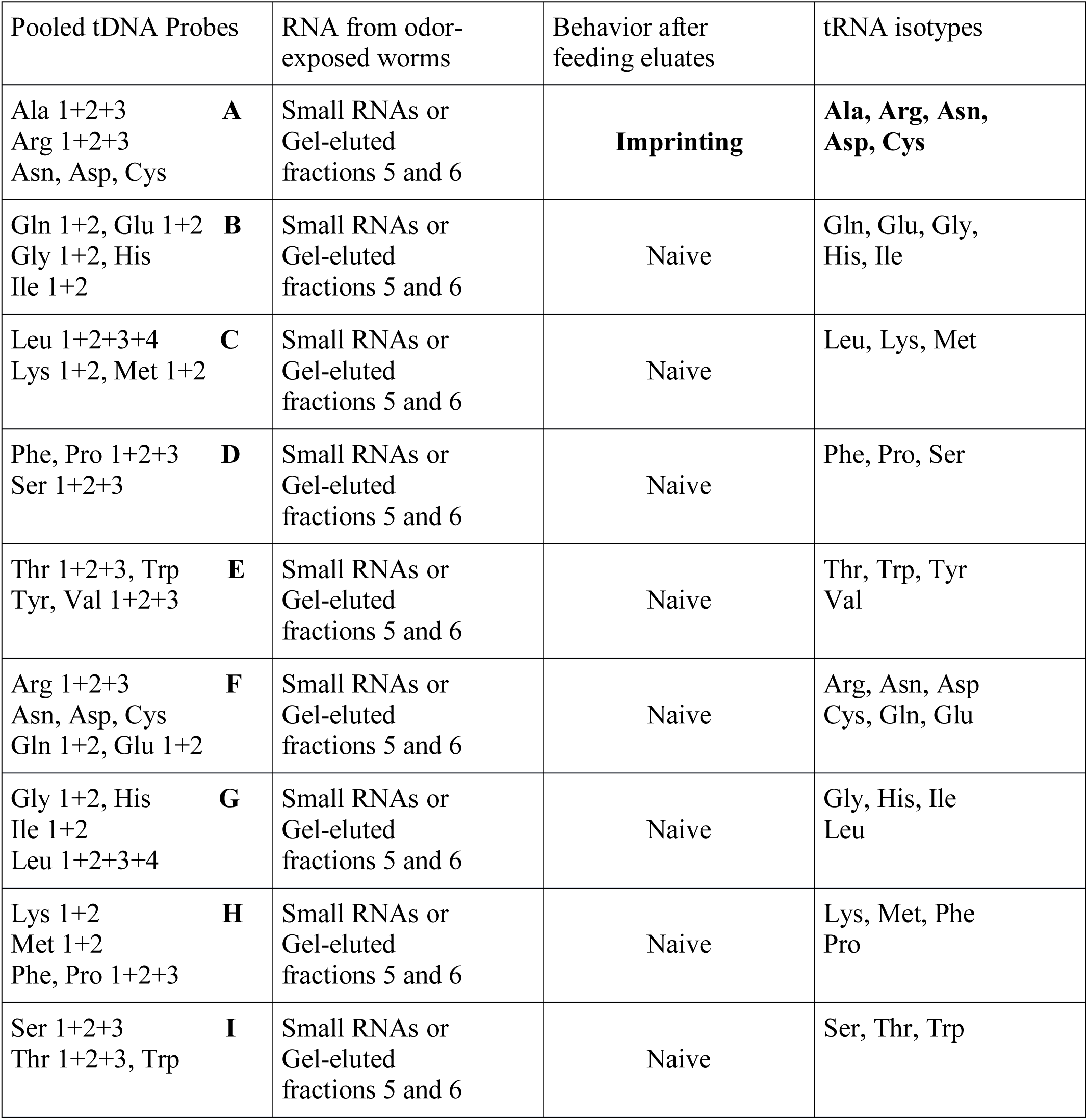
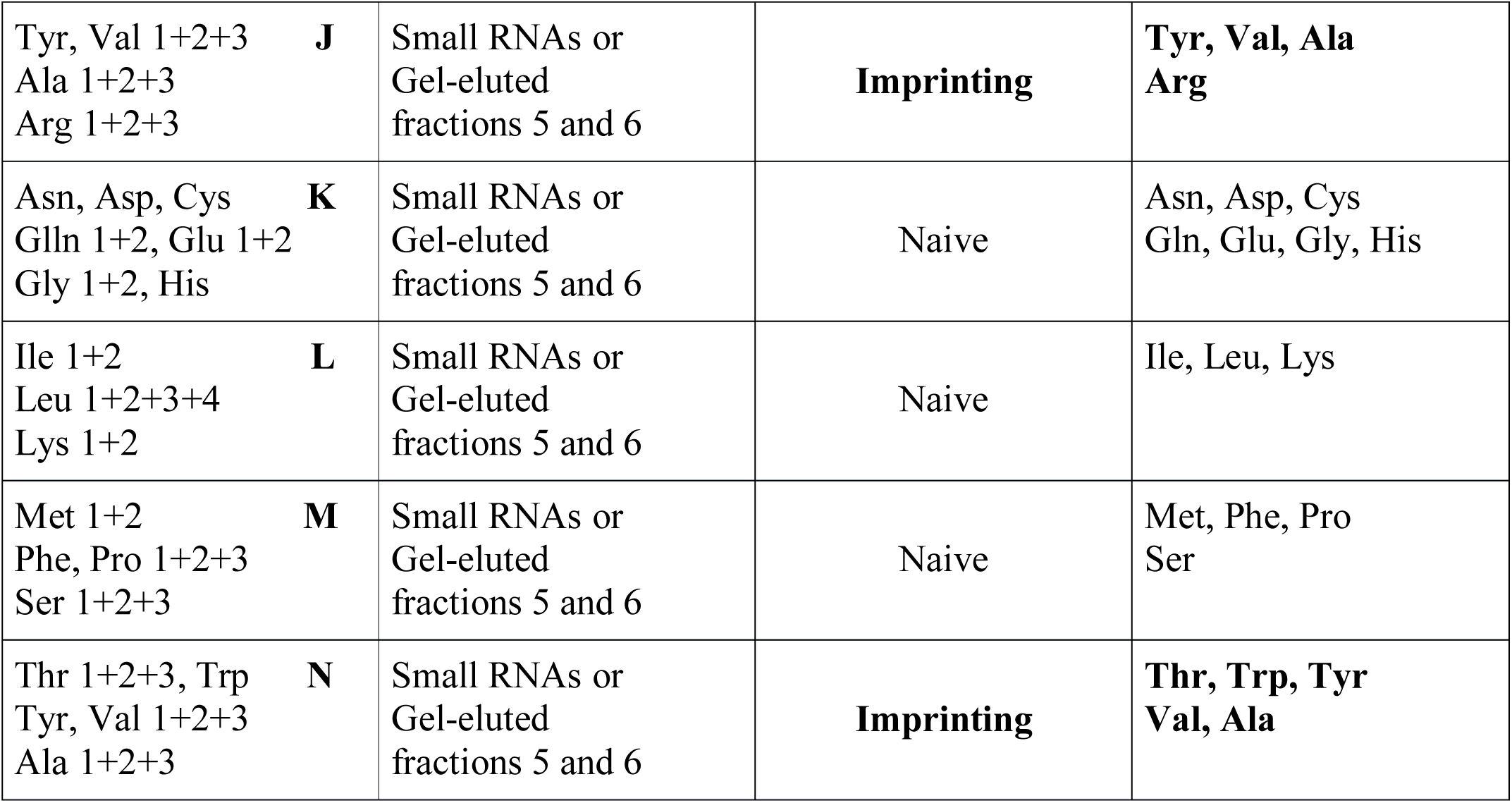
Alanine tRNAs from odor-exposed worms transfer imprinting to naive.

| Pooled tDNA Probes | RNA from odor-exposed worms | Behavior after feeding eluates | tRNA isotypes |
| --- | --- | --- | --- |
| Ala 1+2+3 **A**<br>Arg 1+2+3<br>Asn, Asp, Cys | Small RNAs or Gel-eluted fractions 5 and 6 | **Imprinting** | **Ala, Arg, Asn, Asp, Cys** |
| Gln 1+2, Glu 1+2 **B**<br>Gly 1+2, His<br>Ile 1+2 | Small RNAs or Gel-eluted fractions 5 and 6 | Naive | Gln, Glu, Gly, His, Ile |
| Leu 1+2+3+4 **C**<br>Lys 1+2, Met 1+2 | Small RNAs or Gel-eluted fractions 5 and 6 | Naive | Leu, Lys, Met |
| Phe, Pro 1+2+3 **D**<br>Ser 1+2+3 | Small RNAs or Gel-eluted fractions 5 and 6 | Naive | Phe, Pro, Ser |
| Thr 1+2+3, Trp<br>Tyr, Val 1+2+3 **E** | Small RNAs or Gel-eluted fractions 5 and 6 | Naive | Thr, Trp, Tyr<br>Val |
| Arg 1+2+3 **F**<br>Asn, Asp, Cys<br>Gln 1+2, Glu 1+2 | Small RNAs or Gel-eluted fractions 5 and 6 | Naive | Arg, Asn, Asp<br>Cys, Gln, Glu |
| Gly 1+2, His **G**<br>Ile 1+2<br>Leu 1+2+3+4 | Small RNAs or Gel-eluted fractions 5 and 6 | Naive | Gly, His, Ile<br>Leu |
| Lys 1+2 **H**<br>Met 1+2<br>Phe, Pro 1+2+3 | Small RNAs or Gel-eluted fractions 5 and 6 | Naive | Lys, Met, Phe<br>Pro |
| Ser 1+2+3 **I**<br>Thr 1+2+3, Trp | Small RNAs or Gel-eluted fractions 5 and 6 | Naive | Ser, Thr, Trp |

|  |  |  |  |  |
| --- | --- | --- | --- | --- |
| Tyr, Val 1+2+3<br>Ala 1+2+3<br>Arg 1+2+3 | **J** | Small RNAs or<br>Gel-eluted<br>fractions 5 and 6 | **Imprinting** | **Tyr, Val, Ala**<br>**Arg** |
| Asn, Asp, Cys<br>Gln 1+2, Glu 1+2<br>Gly 1+2, His | **K** | Small RNAs or<br>Gel-eluted<br>fractions 5 and 6 | Naive | Asn, Asp, Cys<br>Gln, Glu, Gly, His |
| Ile 1+2<br>Leu 1+2+3+4<br>Lys 1+2 | **L** | Small RNAs or<br>Gel-eluted<br>fractions 5 and 6 | Naive | Ile, Leu, Lys |
| Met 1+2<br>Phe, Pro 1+2+3<br>Ser 1+2+3 | **M** | Small RNAs or<br>Gel-eluted<br>fractions 5 and 6 | Naive | Met, Phe, Pro<br>Ser |
| Thr 1+2+3, Trp<br>Tyr, Val 1+2+3<br>Ala 1+2+3 | **N** | Small RNAs or<br>Gel-eluted<br>fractions 5 and 6 | **Imprinting** | **Thr, Trp, Tyr**<br>**Val, Ala** |

Northern blot analysis were performed to identify which tRNAs are present in the imprinting positive fractions (***Figure 3b***). We show here the analysis of naive (N3 to N6), CI-exposed (C3 to C6) and BA-exposed (B4 to B7) fractions using probes specific to most tRNA isotypes used in ***Figure 3a***. While tRNA^Leu^ co-migrate with fractions 3, tRNA^Glu^ co-migrate with tRNA^Gln^ mostly in fractions 7, but are also present in fractions 6, tRNA^Lys^ migrate mostly in fractions 6, but are also present in fractions 5. Northern blots support the results obtained using microbeads purification: Alanine tRNAs are mostly present in the imprinting fractions 5 and 6.

The oligonucleotide 1 to 4 described in Methods, purify the three different tRNA^Ala^ isoacceptors used by *C. elegans*, i.e. tRNA^Ala^ (AGC), tRNA^Ala^ (CGC) and tRNA^Ala^ (UGC). To further discriminate between these Alanine tRNA isoacceptors, each purified tRNA was assayed for imprinting transfer.

RNA species eluted from the microbeads bearing the tRNA^Ala^ (AGC) or the tRNA^Ala^ (CGC) probes, do not transfer imprinting. By contrast, the RNA species bound to the two probes corresponding to, respectively, the 5’ and the 3’ halves of the tRNA^Ala^ (UGC) transfer imprinting to naive worms (***Figure 3c)***.

tRNAs eluted from microbeads bearing the or the oligonucleotides, do not transfer imprinting. By contrast, the tRNAs bound to the two oligonucleotides corresponding to, respectively, the 5’ and the 3’ halves of the transfer imprinting to naive worms Our findings suggest that the tRNA^Ala^ (UGC) isodecoder is able to transfer odor-specific imprinting to naive. To further prove this single tRNA molecule is indeed able to carry and transfer several different odor-specific information, we also exposed, besides to BA and to CI, worms to isoamyl alcohol (IA), a third attractive odorant (***Bargmann, 1993***), for which imprinting after early exposure has been et al. demonstrated (***Remy, 2005***).

The use of the microMACS Streptavidin MicroBeads protocol allowed a fast screening of tRNAs activity. We followed the recommended protocol using stringent hybridization conditions and high salt washing buffer. However, to obtain highly purified tRNA^Ala^ (UGC) molecules, visualize the purified tRNA, estimate the purification yields and assess the efficiency of imprinting tranfer, we scaled up our purification by using chromatography on Streptavidin Sepharose beads coupled to the 3’-biotinylated Ala (TGC)-3’ oligonucleotide, as described in Methods.

Starting with 1.10^4^ worms, the amount of eluted tRNA was too low to be visible by ethidium bromide staining after gel migration. Therefore, we used Cy3 labelling, as described in the Methods, to detect the tRNAs after gel fractionation.

As shown on ***Figure 4a***, a major band of tRNA^Ala^ (UGC) from naive and from CI, IA or BA-exposed worms is eluted from the beads. To eliminate any other RNA molecule that may have been co-purified, we cut the bands out the gel, and eluted the tRNAs. The estimated yields were approximately of 1ng pure tRNA^Ala^ (UGC) obtained from 1.10^4^ adult worms. Naive wild-type worms were fed on 1/10 µl of each gel-eluted tRNAs and assayed for CI, IA or BA chemotaxis. Feeding naive worms on gel-eluted tRNA^Ala^ (UGC) from CI-exposed (CI) worms increases chemo-attraction to CI, compared to feeding tRNA^Ala^ (UGC) from naive unexposed (NA) worms. Feeding on tRNA^Ala^ (UGC) from IA or BA-exposed (IA, BA) worms, respectively increases IA or BA responses specifically, compared to feeding tRNA^Ala^ (UGC) from naive (NA) animals (***Figure 4b***).

**Figure 4.**
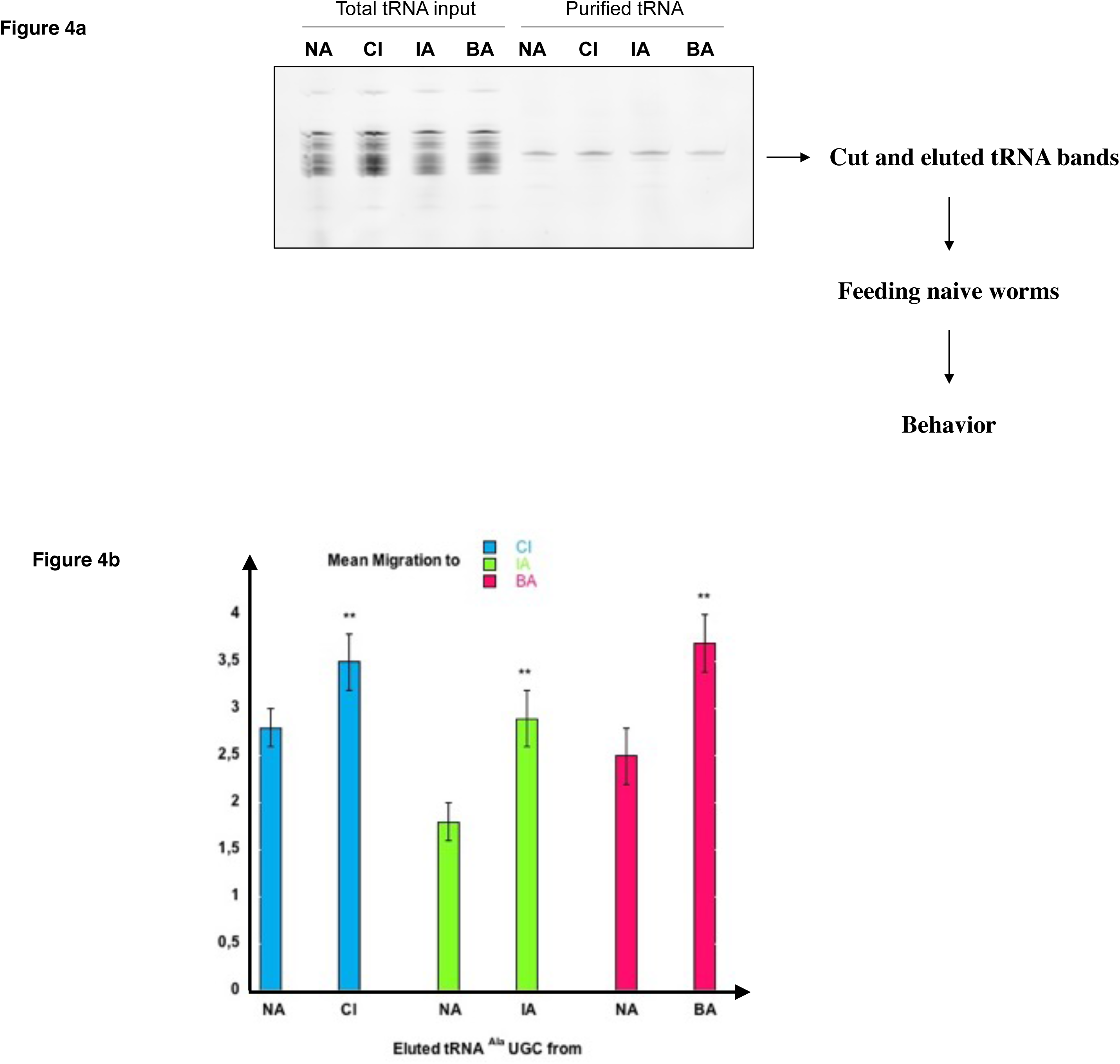
Highly purified tRNA eluted from the tDNA Alanine TGC probe transfers odor-specific imprinting to naive. **(a)** Analysis on 15 % denaturing acrylamide gel of pCp-Cy3 labelled total tRNA and tRNA eluted from 3’-biotinylated tDNA Alanine TGC probe. tRNAs are from naive unexposed (NA), CI-exposed (CI), IA-exposed (IA) or BA-exposed (BA) N2 worms. tRNA bands indicated by the arrow were cut out, eluted, and resuspended in 4 lu H2O to feed naive worms. **(b)** One tenth of each gel-eluted tRNA was added to the food of naive worms. tRNA fed worms were assayed at the adult stage for odor-specific chemotaxis responses (**p-value < 0.01).

These results strongly suggest that odor-stimulated worms produce odor-specific forms of a unique tRNA^Ala^ (UGC) molecule, each of it carrying and transferring an odor-specific information, according to a specific early olfactory experience. Furthermore, imprinting is efficiently transferred by very low amounts of tRNA; we found that 0.001 µl of gel-eluted tRNAs bands (shown on the ***Figure 4a*** gel), containing less than one pg of tRNA^Ala^ (UGC), transfers imprinting to a population of 20 worms.

Enhanced response after odor-exposure is inherited by the F1 unexposed worm generation, while it is lost in the F2 generation. However, olfactory imprinting is stably fixed and stably inherited in worm populations after five worm generations were odor-exposed (***Remy, 2010***).

Imprinting via tRNA^Ala^ (UGC) feeding might be able to recapitulate the inheritance pattern of imprinting after odor-exposure. To assess this question, we fed naive N2 worms for seven successive generations on CI-tRNA^Ala^ (UGC), able to that transfer a CI imprint to naive. As schematically outlined (***Figure 5***), we compared CI responses of the seven CI-tRNA fed generations and of their naive progeny grown without tRNA addition up till the fourth generation. We found that a CI imprint is passed to the first but not to the second naive generation issued from one to five generations of CI-tRNA fed animals. However, CI imprinting is stably maintained in naive generations issued from worms fed on CI-tRNA at least for six successive generations (***Figure 5***). In order to assess the stability of inheritance after the 6th CI-tRNA fed generation, we grew more generations without adding tRNAs. We observed that multi-generationally tRNA-triggered imprinting, as odor-triggered imprinting, is stably maintained in worm progeny. Although it takes six instead of five generations, odor-tRNAs feeding elicit the same long-term stably inherited behavioral change as early odor-exposure.

**Figure 5.**
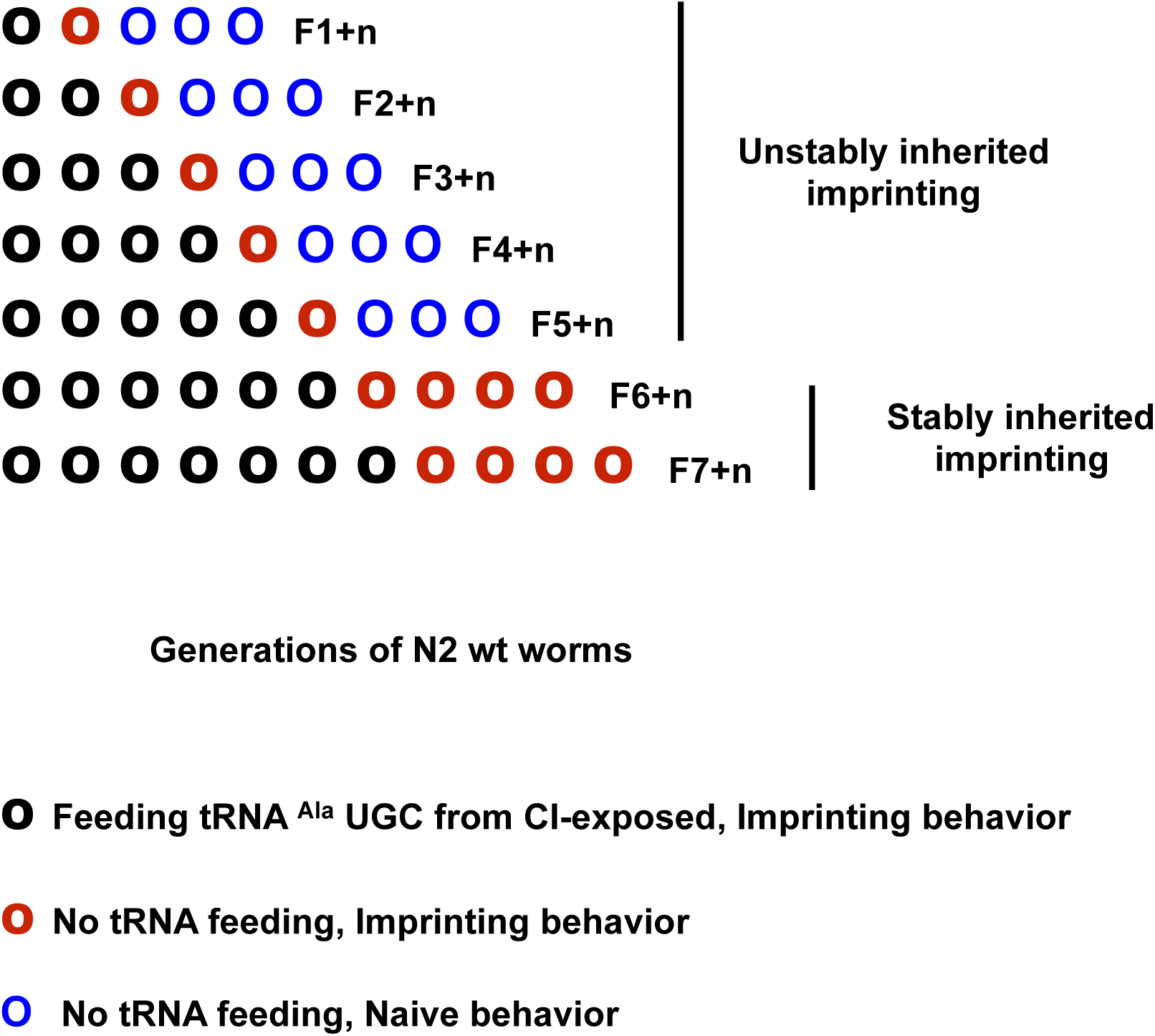
Olfactory imprinting triggered by tRNA^Ala^ (UGC) can be fixed and stably inherited after feeding at least six generations. Imprinting segregation in the progeny of multigenerational imprinting tRNA fed worm populations (schematic representation). CI imprint (CI-tRNA) produced by CI-exposed worms was eluted from microbeads bound to the Alanine tDNA 3’-TGC probe. Naive worm populations were fed on CI-tRNA for one (F1) to seven (F7) successive generations (**O**). Naive generations grown without addition of RNAs were obtained from each of the seven (F1+n to F7+n, here n=4) CI-tRNAs fed populations. CI-tRNAs fed worms passed a CI-imprint to the first naive generation (**O)**. Imprinting is definitely lost at the second naive generation issued from one to five CI-tRNAs fed generations (O O O…); by contrast, it is fixed and definitely maintained in worm populations issued from six and from seven CI-tRNAs fed generations (**O O O O…)**.

tRNA^Ala^ (UGC) from odor-exposed worms are able to stably reprogram odor-specific responses, producing worm populations displaying odor-specific enhanced responses, compared to parental worms.

To alter odor responses and imprint next generations, odor-specific tRNAs added to worm food must enter worm tissues through the intestinal cells. If odor stimulation leads to the synthesis of odor-specific tRNAs within chemo-receptor neurons, then diffusion to germ-line cells would be required for imprinting inheritance.

The tRNAs secondary structure is made of loops joined by double-stranded streches. Uptake and diffusion of double-stranded RNA (dsRNA) support systemic silencing by RNA interference in *C. elegans*. We hypothesized tRNAs could take the paths used by dsRNA.

We studied imprinting and its inheritance in mutant worms bearing amino-acid substitutions in one of two *C. elegans* double-stranded RNA selective transporters SID-1 or SID-2 (***Jose et al., 2009; McEwan et al., 2012)***. The dsRNA selective transporter SID-2 is exclusively localized to the apical membrane of intestinal cells, where it is responsible for the initial binding and internalization of dsRNA from the intestinal lumen (***Marré et al., 2016***). Due to its high expression in germ-line cells, the dsRNA selective importer SID-1 has been involved in transgenerational diffusion of neuronally expressed mobile dsRNAs (***Devanapally et al., 2014***).

Worms with the *sid-1 (qt2),* the *sid-1 (pk3321)* and the *sid-2 (qt13)* mutations show a wild-type imprinting behavior after an early odor exposure (**Table 2**, Odor-exposure). While odor-triggered imprinting was F1 inherited in N2 wild-type and *sid-2 (qt13)* mutant worms, imprinting inheritance was impaired by the two *sid-1 (qt2)* and *sid-1 (pk3321)* substitutions mutations (**Table 2**, Inheritance). Furthermore, while CI-RNA feeding do imprint naive wild-type, *sid-1 (qt2)* and *sid-1 (pk3321)* worms, the *sid-2 (qt13)* mutant worms do not acquire a CI imprint via RNA feeding (**Table 2**, RNA feeding). These results suggest intertissular and intergenerational imprinting tRNAs use the SID-1 and SID-2 dependent paths described for linear dsRNA motility in *C. elegans*.

**Table 2:** tRNAs use the SID-1 and SID-2 dsRNA-specific transporters to, respectively, support imprinting inheritance and imprinting transmission via feeding. Wild-type N2 larvae and larvae with the *sid-1 (qt2), sid-1 (pl3321)* and *sid-2 (qt13)* mutations were either CI 1/300 exposed, fed on 100 ng of RNA from CI 1/300 exposed N2 (CI-RNA), or unexposed as control naive. Mean Migration Indices (MMI) to CI 1/300 of naive worms (A), of CI-exposed worms (B), of unexposed F1 worms from CI-exposed (C), and of CI-RNA fed worms (D), were determined as described (experimental repeats > 4).

| Behavior<br>Genotypes | Chemotaxis of<br>naive unexposed | Imprinting after<br>odor-exposure | Imprinting<br>Inheritance | Imprinting after<br>CI-RNA feeding |
| --- | --- | --- | --- | --- |
| **N2 wt** | **1.6 +/- 0.2** | **2.8 +/- 0.3**<br>**p<0.001** | **2.8 +/- 0.3**<br>**p<0.001** | **2.9 +/- 0.3**<br>**p<0.001** |
| ***sid-1 (qt2)*** | **1.8 +/- 0.3** | **3.2 +/- 0.2**<br>**p<0.001** | **1.7 +/- 0.3**<br>**(= naive)** | **3.0 +/- 0.2**<br>**p<0.001** |
| ***sid-1 (pk3321)*** | **1.8 +/- 0.3** | **2.9 +/- 0.2**<br>**p<0.001** | **1.8 +/- 0.3**<br>**(= naive)** | **2.8 +/- 0.3**<br>**p<0.001** |
| ***sid-2 (qt13)*** | **0.4 +/- 0.2** | **1.5 +/- 0.2**<br>**p<0.001** | **1.6 +/- 0.2**<br>**p<0.001** | **0.5 +/- 0.2**<br>**(= naive)** |
|  | **A) Naive**<br>**CI MMI** | **B) CI-exposed**<br>**CI MMI** | **C) Unexposed F1**<br>**generation**<br>**CI MMI** | **D) CI-RNA fed**<br>**CI MMI** |

*A*ttractive odors, when present in the first *C. elegans* larval stage environment, would trigger the production of different forms of tRNA^Ala^ (UGC), each bearing an odor-specific signature. The technology available to analyse RNA chemical modifications is making progress and rapidly developing. These methods, either based on biochemical or sequencing approaches, mostly apply however to known identified single modifications. To this day, there is no straight forward available method able to accurately describe the whole quantitative and qualitative pattern of bases modifications present on a single purified RNA molecule (***Schaefer et al., 2017; Jonkhout et al., 2017; Motorin and Helm, 2019*)**.

tRNAs bases are extensively modified. The modified nucleotides combinations potentially produce a great number of chemically different tRNA molecules. How such combinatorial chemical diversity is produced, controlled and linked to tRNA biological functions remains to be described. We hypothesize that differential combinations of bases modifications would form the odor-specific codes carried by tRNA^Ala^ (UGC) after odor stimuli.

Aiming to identify some differences between naive and odor induced tRNAs, we based our investigations within the scope of the currently limited knowledge on tRNA modifications in *C. elegans*. Most tRNA-modifying enzymes and biosynthesis pathways are described in the yeast, but remain unknown in the worm.

Due to the essential role of tRNAs in protein translation, the most studied tRNA chemical modifications have been those affecting the wobble position 34, the first base of the anticodon, in particlular Uridine 34.

Wobble U modifications are considered of critical functional importance in codon-anticodon recognition as they might improve tRNA aminoacylation kinetics and prevent translational frame-shifts (***Larsen et al., 2015; Nedialkova and Leidel, 2015; Deng et al., 2015***).

tRNA^Ala^ (UGC) indeed carries an uridine at position 34, by contrast to the two other Alanine tRNA (AGC) and tRNA (GCG). Several U34 modifications require the activity of the tRNA modifying Elongator complex (***Karlsborn et al., 2014***).

The yeast and mammalian Elongator complexes are composed of two copies of two sub-complexes associating of the three ELP-1-2-3 and of the three ELP-4-5-6 subunits (***Glatt et al., 2012, Dauden et al., 2017***). However, the *C. elegans* elongator complex, ELPC, may function as a 1-2-3 sub-complex, since the elongator sub-units 5 and 6 genes are absent from the worm genome.

Recent structural and functional analysis of tRNAs processing by the yeast elongator complex indicates that the carboxymethylation (cm5) of 11 tRNAs carrying U34, including the tRNA^Ala^ (UGC), requires the activity of the catalytic 1-2-3 sub-complex (***Dauden et al., 2019***). The cm5 modification will further lead to the formation of 5-methoxycarbonylmethylated (mcm5), 5-methoxycarbonylmethyl-2-thiolated (mcm5S2) and 5-carbamoylmethylated (ncm5) modified uridines. In the yeast, ELP-1 and ELP-3 sub-units both interact with tRNAs. The Methanocaldococcus infernus archeon, which express none but the ELP-3 sub-unit, do performs the U34 carboxymethylation reaction. It has been shown that the *C. elegans* elongator complex also controls the synthesis of the mcm5, mcm5S2 and ncm5 modifications of tRNA uridines (***Chen et al., 2009***).

Elongator has been implcated in a great variety of nuclear and cytoplasmic cellular functions, including transcription elongation, chromatin remodeling, exocytosis, zygotic paternal DNA demethylation, and neuronal development (***Creppe et al., 2009; Okada et al., 2010; Solinger et al., 2010; Tielens et al., 2016; Dalwadi and Yip, 2018)***. The multifunctionality of Elongator could be partly explained by structural analysis (***Dauden et al., 2017, 2019).*** ELP3 contains a C-terminal Lysine Acetyl Transferase (KAT) domain and a domain with sequence homology with the S-adenosylmethionine (SAM) domain. EPL-3 KAT could acetylate a number of targets, including histones and neuronal alpha-tubulin (***Creppe et al., 2009; Solinger et al., 2010).*** In the mouse, the radical SAM domain but not the KAT domain has been involved in paternal genome demethylation, suggesting the two ELP3 functional domains could play different roles (***Okada et al., 2010***). Based on over 80% primary sequence identity, ELPC-3, the worm Elongator sub-unit 3, may carry the same functional domains described for yeast and mammals.

Recent studies reported the effects of mutations inactivating the *C. elegans* Elongator sub-units. It is important to notice that worms harboring mutations in either *elpc-1, elpc-2* or *elpc-3* genes display neuronal and behavioral phenotypes (***Chen et al., 2009; Solinger et al., 2010; Kawamura and Maruyama, 2019***). This suggests a remarkable evolutionary conservation of the elongator functions in neurodevelopment ***(Creppe, 2009; Kojic and Wainwright, 2016)***.

To see if ELPC-3 could be involved in the regulation of chemo-attraction, we used worms carrying different chromosomal deletions of the *elpc-3* gene ***(Figure 6)***. The 1503 bp *ok2452* deletion span the whole putative ELPC-3 Radical SAM domain, mapped by homology between aminoacids 89 and 300, while the 355 bp *tm3120* deletion may leave it intact. We mapped the putative C-terminal ELPC-3 KAT domain between aminoacids 436 and 547, based on sequence homology with the yeast and mammalian ELP-3 KAT domains. Using the CRISP-Cas9 technology, we produced worms with a 375 bp deletion, called here *del-KAT*, which eliminates the KAT domain from ELPC-3 ***(Figure 6a)***. We found that, thus to variable extents, all tested *elpc-3* deletion mutants display significantly reduced chemo-attractive responses to CI, BA and IA, compared to wild-type N2 worms (***Figure 6b***).

**Figure 6.**
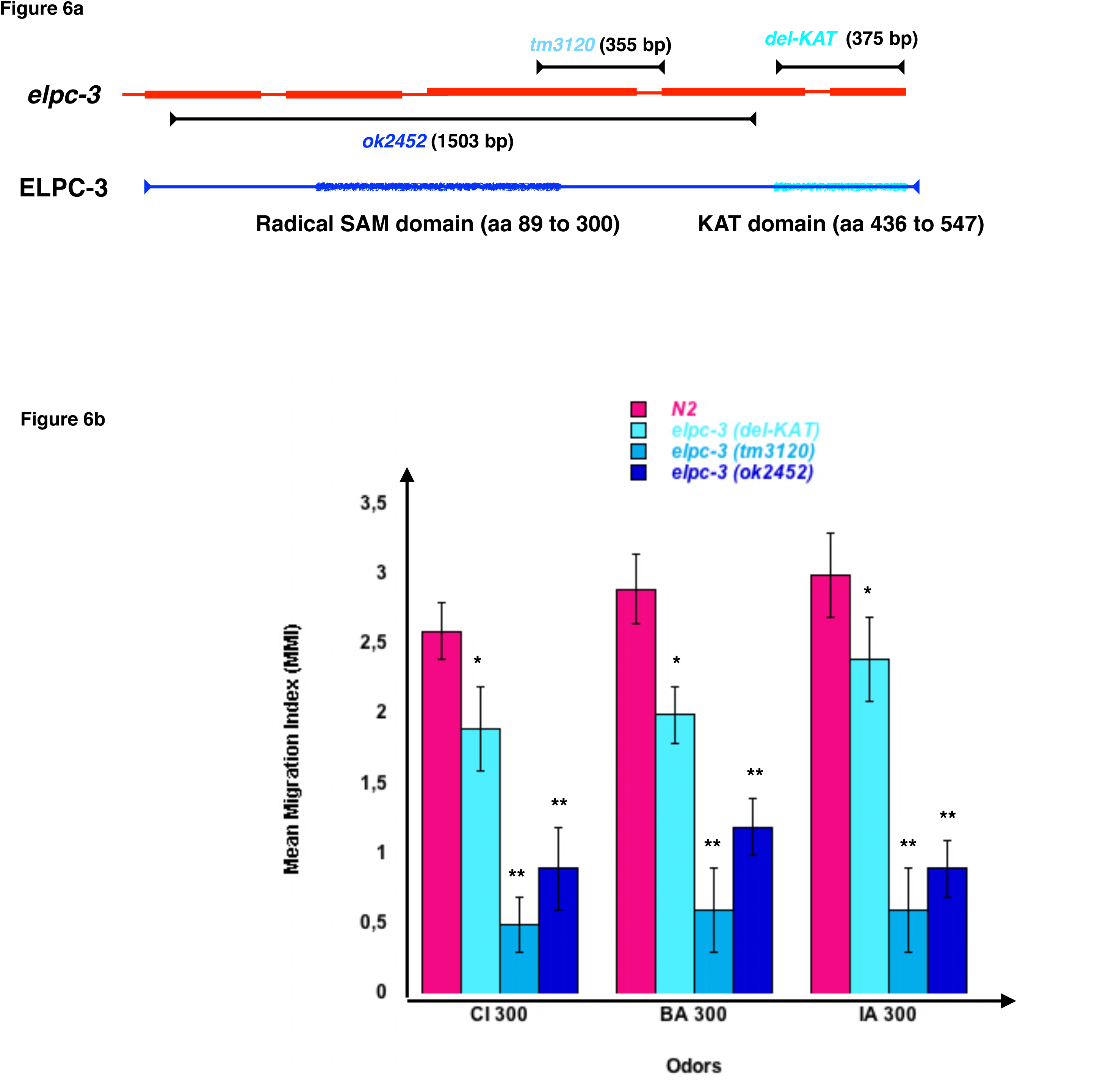
*C. elegans* chemo-attractive behavior requires a functional elongator complexe sub-unit 3 (ELPC-3). **(a)** Schematic representation of the *C. elegans* Elongator sub-unit 3 (*elpc-3*) gene, *elpc-3* gene deletion, and of the putative Radical SAM and Lysine Acetyl Transferase KAT domains of the ELPC-3 protein. **(b)** Chemo-attractive behavior of worms bearing different deletion of the *elpc-3* gene, compared to wild-type N2. Mean Migration Indices (MMI) to Citronellol (CI 300), Benzaldehyde (BA 300) and Isoamyl alcohol (IA 300) were determined as described for 4 days old worms (experimental repeats ≥ 4, *p-value < 0.05, **p-value < 0.01).

We next tested the behavioral effects of the three different *elpc-1* gene deletions described in ***Figure 7a***. Worms bearing the *ng10* 2050 bp deletion could not produce ELPC-1, while worms with the *tm2149* 275 bp and the *tm11684* 95 bp *elpc-1* deletions could produce C-terminal truncated forms of ELPC-1. By contrast to the *elpc-3* mutants, chemo-attraction is not significantly affected by the *elpc-1* gene deletions, compared to wt N2 (***Figure 7b)***.

**Figure 7.**
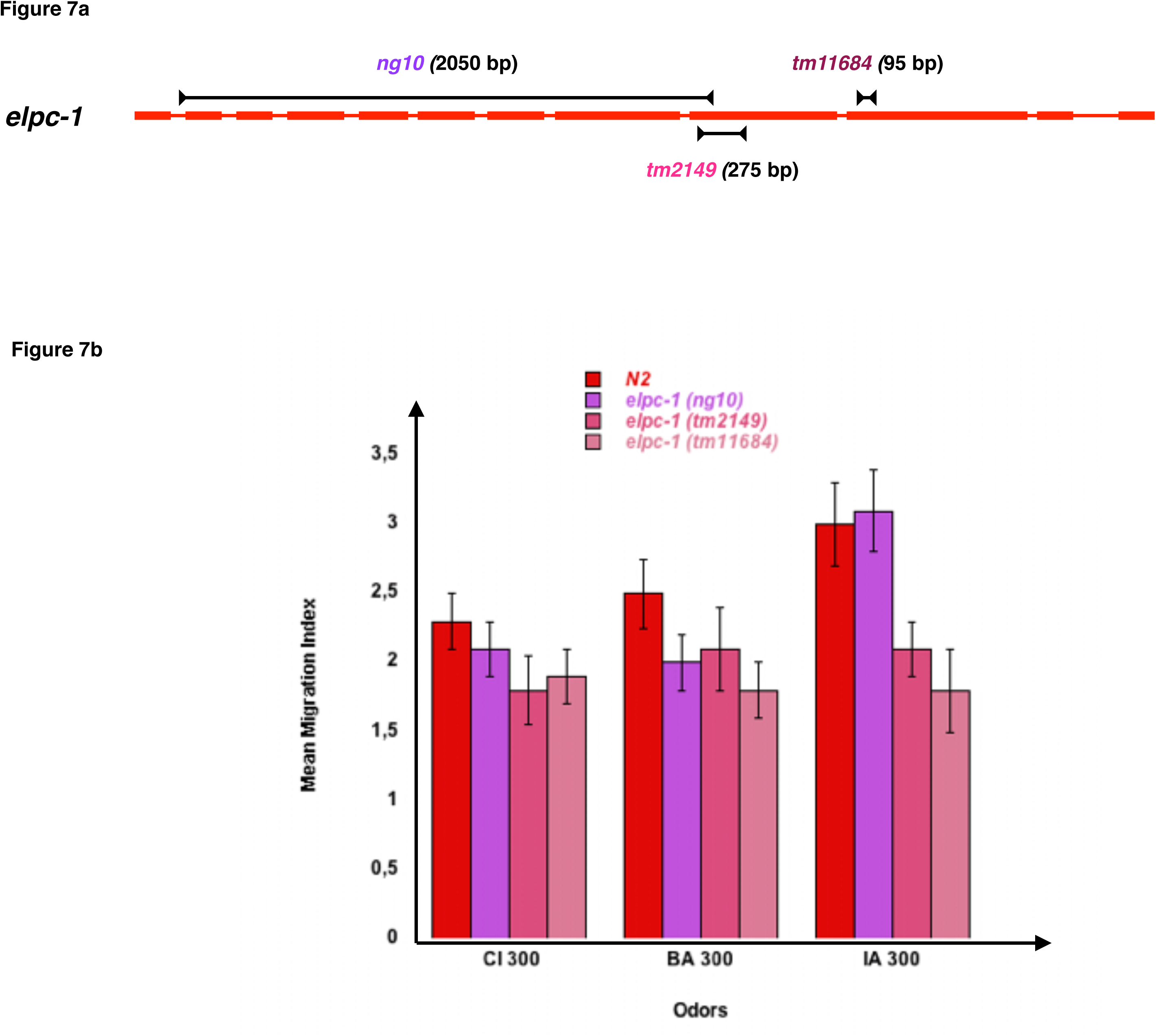
Inactivating the *C. elegans* elongator complexe sub-unit 1 (ELPC-1) does not significantly modify chemo-attractive responses. **(a)** Schematic representation of the *C. elegans* Elongator sub-unit 1 gene, and of the *elpc-1* gene deletions. **(b)** Chemo-attractive behavior of worms bearing three different deletion of the *elpc-1* gene, compared to wild-type N2. Mean Migration Indices (MMI) to Citronellol (CI 300), Benzaldehyde (BA 300) and Isoamyl alcohol (IA 300) were determined as described for 4 days old worms (experimental repeats ≥ 4).

We reasoned that if the chemotaxis defects of *elpc-3* worms are due to impaired modifications of the tRNA bases, then providing wild-type tRNA through feeding might rescue the behavioral phenotype. We cut and eluted the naive wild-type (NA) tRNAs co-migrating fractions 2 to 7 from the gel showed on ***Figure 2a***.

After feeding *elpc-3 (tm3120)* worms on the pooled 2 to 7 fractions and on each fraction, separately, we found that the whole population of naive tRNA (2-7) and the Alanine tRNAs containing fractions 5 and 6 (N_5_ and N_6_ on ***Figure 3***) indeed restored CI chemotaxis ***(Figure 8a)***. We next asked if feeding the microbead-purified wild-type naive tRNA^Ala^ (UGC) is enough to restore the chemo-attractive defects of *elpc-3* mutants. As shown in ***Figure 8b***, *elpc-3 (tm3120)* and *elpc-3 (2452)* worms indeed recover a wild-type behavior after being fed wild-type tRNA^Ala^ (UGC).

**Figure 8.**
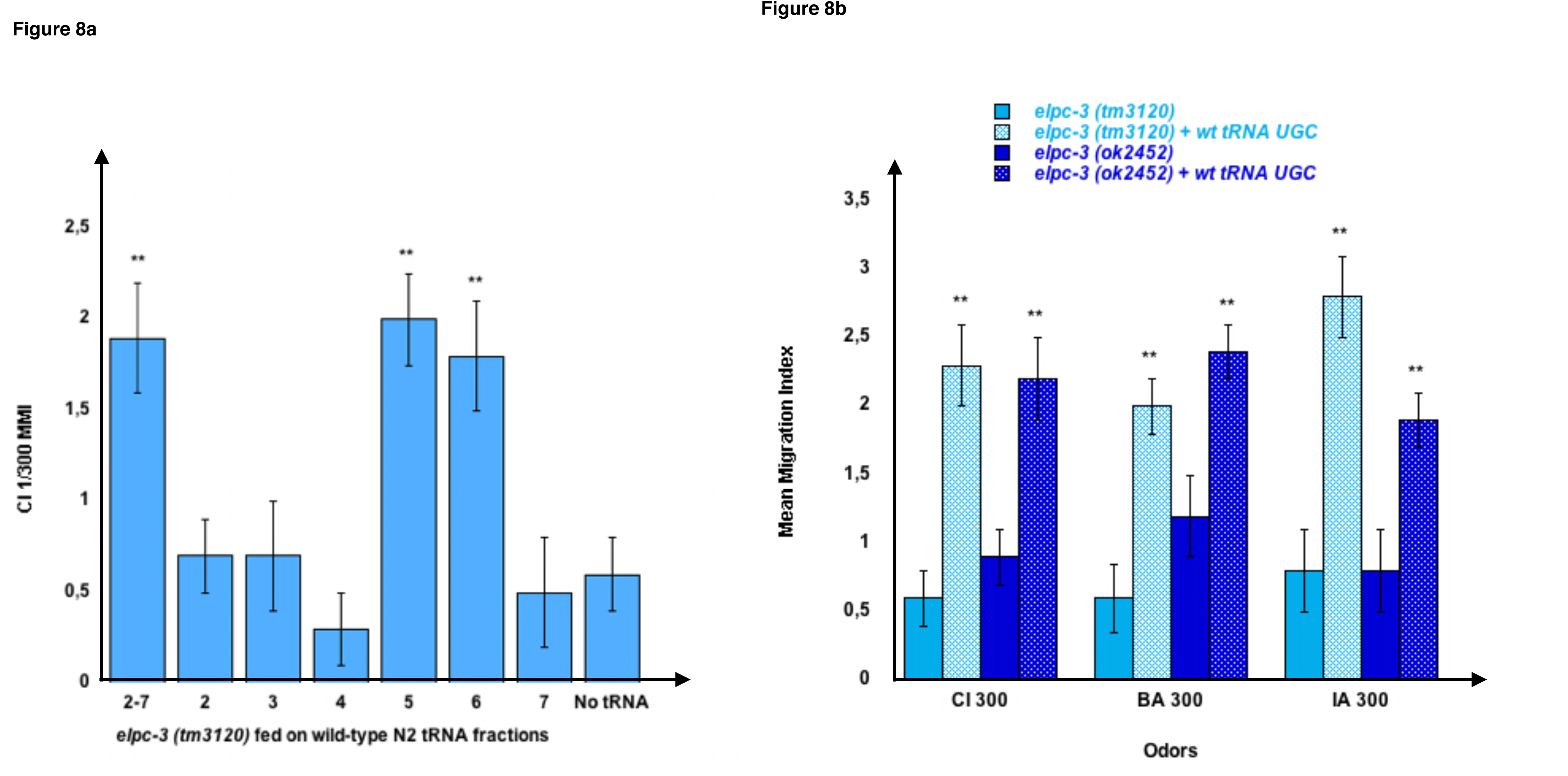
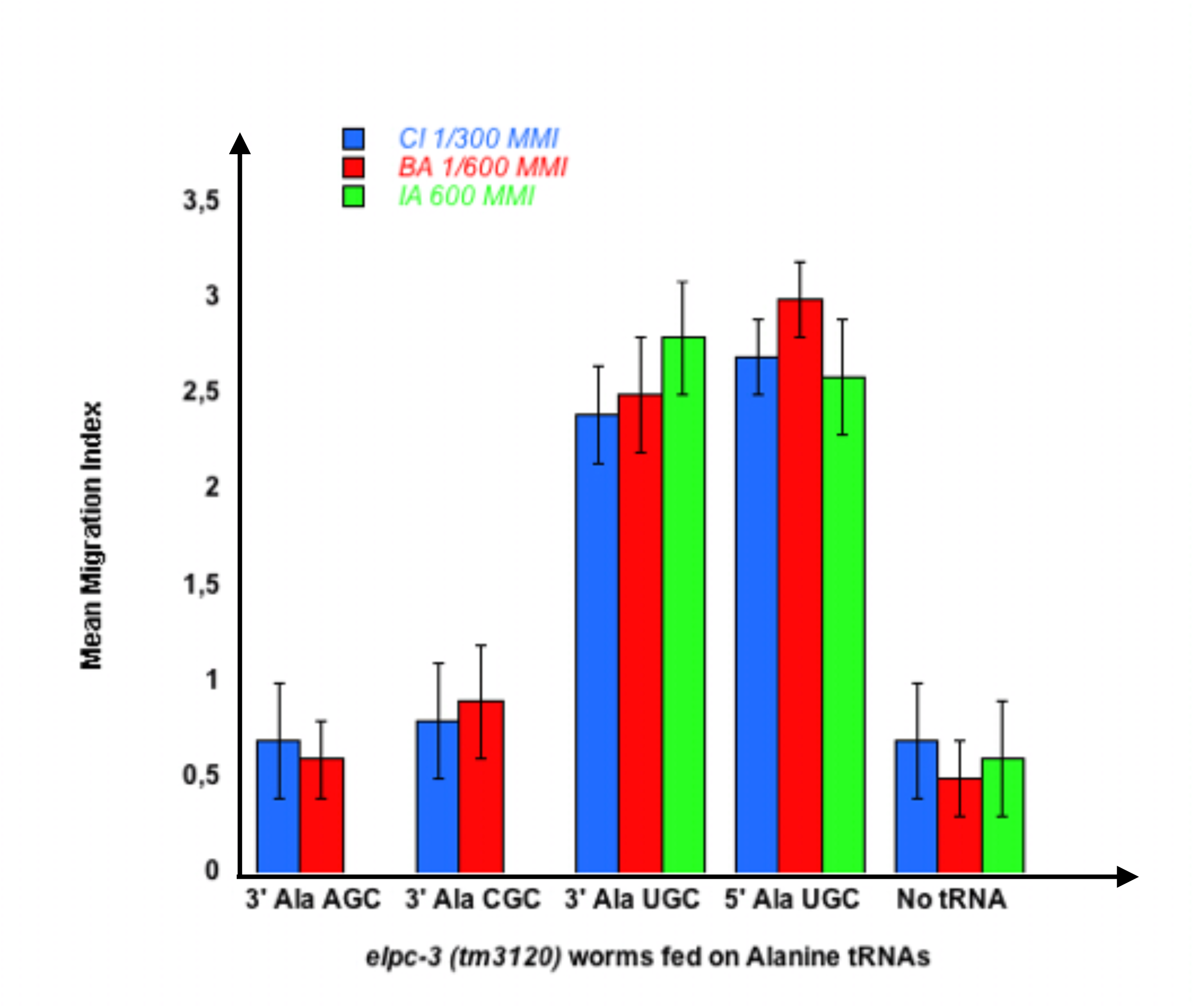
Feeding wild-type tRNA^Ala^UGC fully restores chemo-attraction in *elpc-3* mutants. **(a)** Mean migration indices of *elpc-3 (tm3120)* mutants after they were either unfed (No tRNA), fed on the pooled N2 tRNA fractions (2-7) or fed on individual N2 tRNA fractions (1 to 7). tRNA fractions were gel-extracted as shown on Figure 2. **(b)** Chemo-attractive responses of naive or of wild-type tRNA^Ala^UGC fed *elpc-3 (tm3120)* and *elpc-3 (ok2452)* mutant worms. Mean Migration Indices (MMI) in gradients of Citronellol (CI), Benzaldehyde (BA) and Isoamyl alcohol (IA) at the indicated dilutions, were determined as described (experimental repeats ≥ 4, **p-value < 0.01). **(c)** tRNA^Ala^UGC, but not tRNA^Ala^AGC nor tRNA^Ala^CGC feeding rescues the chemotaxis defects of *elpc-3 (tm3120)* worms.

We next demonstrated that none of the other Alanine tRNAs rescue the *elpc-3* behavioral defects. As shown on ***Figure 8c***, the tRNA^Ala^ (UGC), but not the tRNA^Ala^ (AGC) nor the tRNA^Ala^ (CGC) fully rescues the chemo-attractive defects of *elpc-3 (tm3120)* mutants ***(Figure 8c)***.

We did not observe significant effects of *elpc-1* deletions on chemo-attraction ***(Figure 7b).*** Moreover, feeding wild-type naive tRNA^Ala^ (UGC) modify neither the behavior of *elpc-1 (tm2149)* nor the behavior of *elpc-1 (tm11684)* mutants ***(Supplemental Figure 2)***.

*elpc-3* mutants provided via feeding with the wild-type tRNA^Ala^ (UGC) acquire a wild-type behavior. The rescue of *elpc-3* phenotype by wt tRNA^Ala^ (UGC) suggests *elpc-3* worms may produce a « defective » form of tRNA^Ala^ (UGC), presumably with an altered chemical composition, unable to support the development of chemo-attraction.

We then hypothesized that if the development of a wild-type behavior requires the wt tRNA^Ala^ (UGC) chemical composition, then feeding the *elpc-3* « defective » tRNA^Ala^ (UGC), could transfer the *elpc-3* behavioral phenotype to wild-type. Furthermore, the *elpc-1* mutants, displaying a wild-type chemo-attraction, would produce the« permissive » wild-type chemical form of tRNA^Ala^ (UGC), able to rescue the *elpc-3* behavioral phenotype.

We therefore purified tRNA^Ala^ (UGC) from *elpc-3* and from *elpc-1* worms to assess their behavioral effects. As shown on ***Figure 9a,*** feeding wild-type N2 worms on, respectively, tRNA^Ala^ (UGC) purified from either *elpc-3 (tm3120)*, *elpc-3 (ok2452)* or *elpc-3 (del-KAT)* worms, indeed significantly impairs CI 300 and IA 300 attraction.

**Figure 9.**
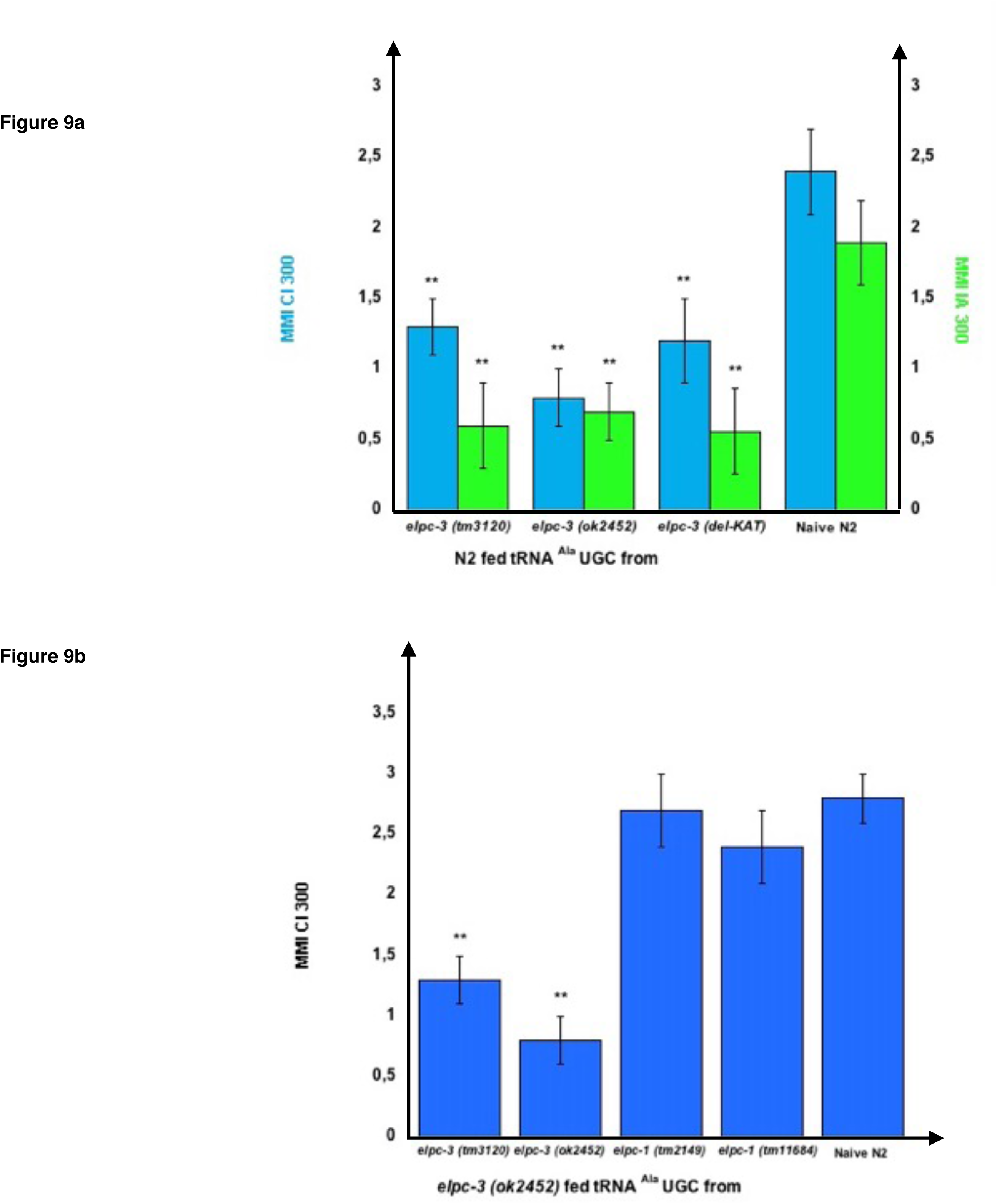
tRNA^Ala^UGC produced by *elpc* mutants transfers mutant phenotypes to wild-type via feeding. **(a)** tRNA^Ala^UGC from the *elpc-3 (tm3120),* from the *elpc-3 (ok2452)* and from the KAT domain deleted *elpc-3 (del-KAT)* mutant worms were microbead-purified as described. Wild-type N2 worms were grown during their whole development in the presence of the indicated *elpc-3* mutants tRNA^Ala^UGC or without tRNA for Naive N2. Mean Migration Indices (MMI) in gradients of Citronellol (CI 300) or Isoamyl alcohol (IA 300) were determined at the adult stage as described (experimental repeats ≥ 4, **p-value < 0.01). **(b)** Feeding tRNA^Ala^UGC from *elpc-1* mutants fully rescues the chemo-attractive defects of *elpc-3* mutants. tRNA^Ala^UGC from *elpc-3 (tm3120), elpc-3 (ok2452), elpc-1 (tm2149),* and from *elpc-1 (tm11684)* mutant worms were microbead-purified as described. *elpc-3 (ok2452)* worms were grown during their whole development in the presence of the indicated *elpc-3* or *elpc-1* mutants tRNA^Ala^UGC. Mean Migration Indices (MMI) of tRNA fed *elpc-3 (ok2452)* and of Naive N2 wild-type in Citronellol (CI 300) gradients were determined at the adult stage as described (experimental repeats ≥ 4, **p-value < 0.01).

Furthermore, feeding tRNA^Ala^ (UGC) purified from either *elpc-1 (tm2149)* or *elpc-1 (tm11684)* worms fully rescues the CI 300 chemo-attractive defect of *elpc-3 (ok2452)* worms, while feeding its own tRNA^Ala^ (UGC) or the *elpc-3 (tm3120)* tRNA^Ala^ (UGC) has no rescuing effect ***(Figure 9b)***.

Importantly, the *elpc-3* rescuing effects of wt or *elpc-1* tRNA^Ala^ (UGC) ***(Figure 8a),*** or the transfer of an *elpc-3* mutant phenotype to wt worms by *elpc-3* tRNA^Ala^ (UGC) ***(Figure 9a and b)*** do not last beyond the tRNA fed worm generation.

**Figure 10.**
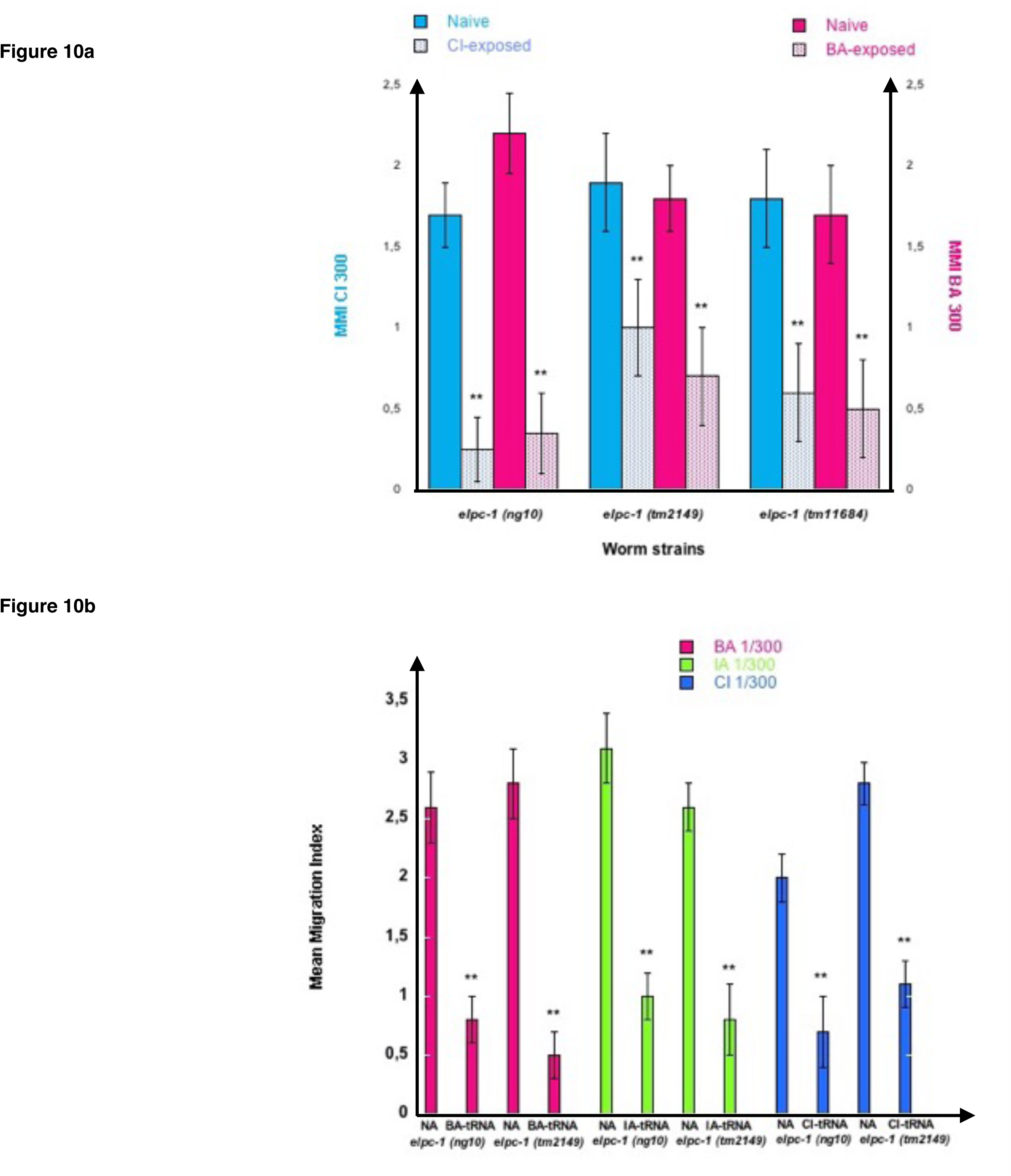
Odor-exposure or feeding odor-tRNA^Ala^UGC strongly reduce odor-specific chemo-attraction in *elpc-1* mutants. **(a)** *elpc-1 (ng10)*, *elpc-1 (tm2149)* or *elpc-1 (tm11684)* mutant worms were either exposed to CI 1/300, to BA 1/300, or unexposed for control Naive. Mean Migration Indices (MMI) of Naive unexposed, of CI-exposed and of BA-exposed *elpc-1* worms in, respectively, CI 1/300 and BA1/300 gradients were determined at the adult stage as described (experimental repeats ≥ 4, **p-value < 0.01). **(b)** *elpc-1 (ng10)* or *elpc-1 (tm2149)* worms were fed (or unfed for control naive NA) on BA-tRNA^Ala^UGC (BA-tRNA), IA-tRNA^Ala^UGC (IA-tRNA) or CI-tRNA^Ala^UGC (CI-tRNA) obtained respectively from BA, IA or CI-exposed wild-type worms. BA, IA or CI-tRNA^Ala^UGC were obtained after microbead purification, gel-separation and gel-elution as described in Figure 4. Mean Migration Indices of Naive unfed (NA) worms were compared to Mean Migration Indices of odor-tRNA fed worms in the respective odor gradients (experimental repeats ≥ 4, **p-value < 0.01).

The *elpc-1* deletions do not affect chemo-attraction ***(Figure 7b)***. We assessed olfactory imprinting in these mutants. In contrast with wild-type, early odor-exposure or odor-specific tRNA^Ala^ (UGC) feeding, do not increase but stably decrease odor responses in *elpc-1* mutants. CI or BA-exposed *elpc-1 (ng10), elpc-1 (tm2149) or elpc-1 (tm11684)* worms display a significant reduction of CI or BA chemo-attractive responses, compared to naive unexposed ***(Figure 10a)***.

Attraction to BA, IA, or CI, significantly decreased after *elpc-1 (ng10)* and *elpc-1 (tm2149)* mutants were fed on, respectively, highly purified by gel elution (as shown on ***Figure 4***), wild-type BA, IA or CI-tRNA^Ala^ (UGC) (***Figure 10b)***.

Thus, in the absence of a functional ELPC-1, olfactory imprints have a negative impact on future adult chemoattraction, as opposed to wild-type.

We furthermore demonstrated that worms can be negatively imprinted by one or several odors, as negative imprinting is stably inherited over further generations. As shown on ***Figure 11***, we submitted all three *elpc-1* deletion mutants to sequential odor-exposures. One generation was exposed to a single odorant, the next generation was (or not) exposed to a second odorant, and the third generation exposed (or not) to a third odorant. IA-exposed worms remain attracted to BA as naive, while sequentially exposed worms to CI then to BA (CI BA) or to CI then to BA then to IA (CI BA IA), lost BA attraction (left part of ***Figure 11***). In the same manner, IA response was lost by IA-exposed and by CI BA IA-exposed *elpc-1* worms, while it remained unaffected in CI BA exposed, compared to naive (right part of ***Figure 11***). The inheritance of odor-specific lowered attraction to either a single, two, or to three different odorants was assessed up to the tenth generation. Altogether, our data clearly indicate that the elongator complex is essential for the development and the regulation of *C. elegans* chemo-attractive behavior.

**Figure 11.**
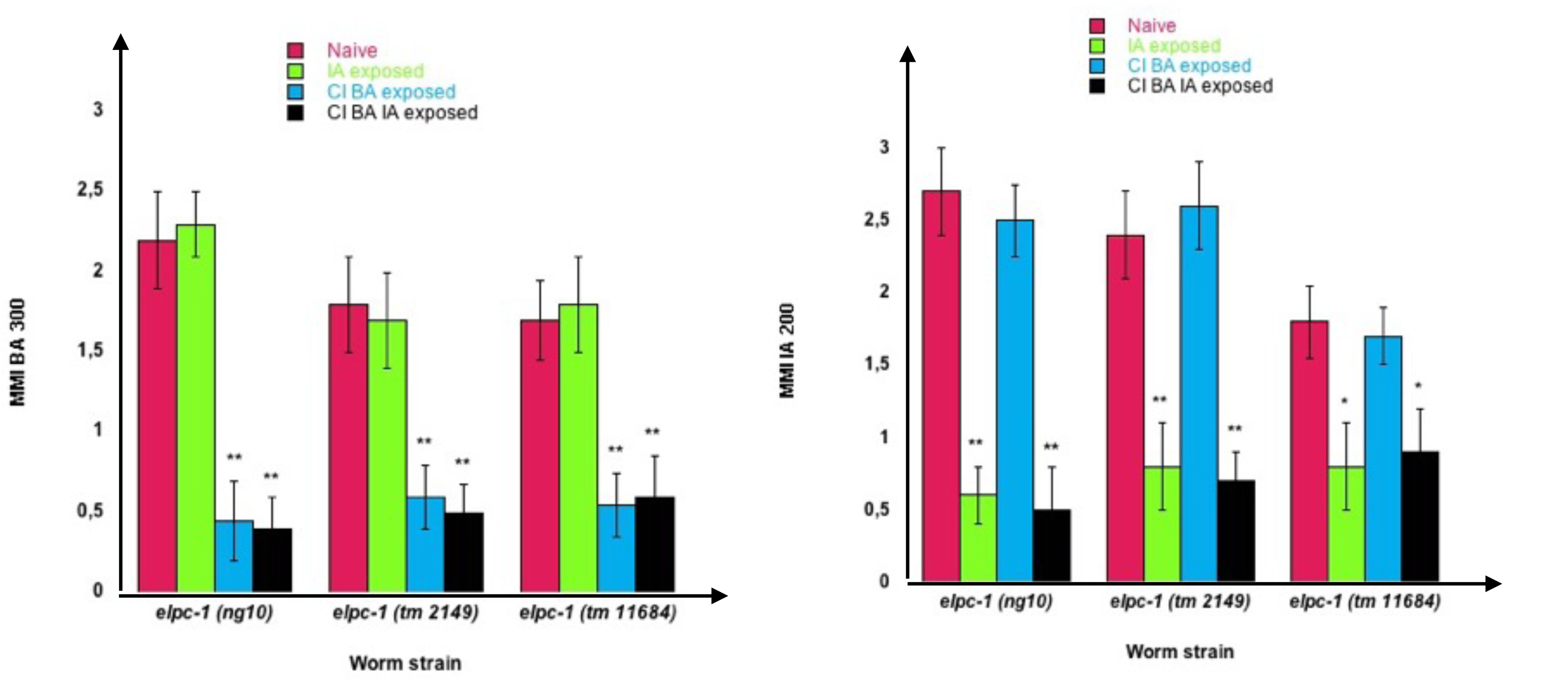
Stable reduction of three odor-specific responses after sequential odor-exposure of *elpc-1* mutants. **(a)** Mean migration Indices (MMI) of Naive unexposed, of IA-exposed, CI BA-exposed and CI BA IA-exposed *elpc-1* worms in BA 1/300 gradients were determined at the adult stage, as described (experimental repeats ≥ 4, **p-value < 0.01) **(b)** Mean migration Indices (MMI) of Naive unexposed, of IA-exposed, CI BA-exposed and CI BA IA-exposed *elpc-1* worms in IA 200 gradients were determined at the adult stage, as described (experimental repeats ≥ 4, **p-value < 0.01).

It has been reported that the total tRNA populations extracted from *elpc-3 (tm3120)* and from *elpc-1 (2149)* lack the mcm5S2U and the ncm5U modifications, and instead contain S2U, as opposed to wild-type tRNAs which carry both mcm5S2U and ncm5U but do not contain S2U (***Chen et al., 2009***). It is believed that these modifications would mainly affect codon-anticodon pairing for the three tRNAs with U at positions 34 and 35, namely Lys (UUU), Glu (UUC) and Gln (UUG). Even if proven true, this simple picture appears far too simple to account for our observations for several reasons:

– There is no specific information regarding if and how U34 are actually modified in the eleven U34 containing tRNAs, that includes Ala (UGC);
– These uridine modifications could be derived from other and can originate other chemical forms of uridine. For instance, mcm5S2U results from the addition of S2U to mcm5U, and ncm5U leads to the formation of at least the three modified uridines mcm5Um, nchm5U or ncm5S2U (***Boccaletto P. et al., 2018***);
– We showed that *elpc-3* and *elpc-1* mutants display different behavioral phenotypes, producing tRNA^Ala^ (UGC) with different activities;

To see if any of the three U modifications could be involved in chemo-attraction or in the positive or the negative imprinting, we used synthetic S2U, mcm5S2U and ncm5U. We first analysed the behavior of the *tuc-1 (tm1297)* mutants, which lack the tRNA sulfurtransferase activity of the TUT-1 enzyme, thus do not synthesize S2U, nor mcm5S2U. Chemo-attraction is impaired by the *tuc-1 (tm1297)* mutation, as shown for CI 300 chemotaxis (***Figure 12a***). We observed a concentration-dependent restoration of CI 300 response after adding increasing amounts of S2U from 2 to 400, to worm food. By contrast, the chemo-attractive defect of the *elpc-3 (tm3120)*⎧M mutants was not restored after feeding the same amounts of S2U, as shown on ***Figure 12a.*** Adding S2U does not alter chemo-attraction in wild-type worms (results not shown).

**Figure 12.**
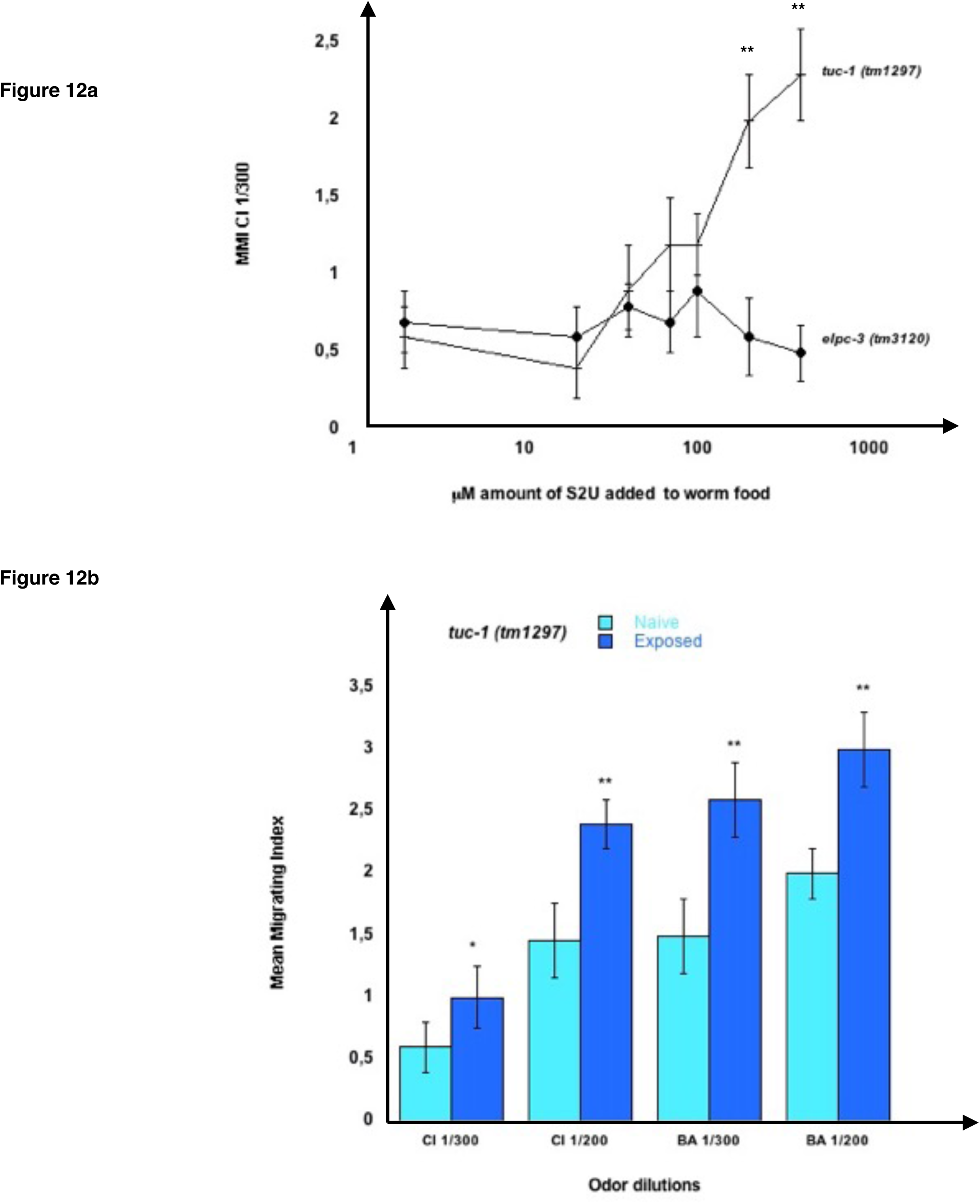
Chemo-attractive behavior of the *tuc-1 (tm1297)* mutant worms. **(a)** Mean migration indices of *tuc-1 (tm1297)* and *elpc-3 (tm3120)* mutant worms in a CI 1/300 gradient in the presence of increasing concentrations of 2-thiouridine (S2U) from 2 to 400 μM. MMI were determined at the adult stage, as described (experimental repeats ≥ 4, **p-value < 0.01). **(b)** *tuc-1 (tm1297)* worms were unexposed for Naive or Exposed to, respectively, CI 1/300, CI 1/200, BA 1/300 or to BA 1/200. Mean Migration Indices in the respective odor-dilutions gradients were determined at the adult stage, as described (experimental repeats ≥ 4, *p-value < 0.05, **p-value < 0.01).

We next assessed imprinting in the *tuc-1 (tm1297)* mutants, and found that, despite a lack of S2U and mcm5S2U, and low chemotaxis responses, these worms positively imprint CI 300, CI200, BA300 and BA 200 (***Figure 12b***).

These results indicate that a behavioral phenotype linked to the absence of a single identified modified nucleotide (here S2U), can be fully rescued via addition of the missing nucleotide in worms environment.

Wild-type tRNAs are believed to have both mcm5S2U and ncm5U, while *elpc* mutants do not. Since imprinting occurs without S2U nor mcm5S2U, as suggested by the behavior of the *tuc-1* mutants, then imprinting could, at least partially, involve ncm5U. In order to test this hypothesis, we fed wt, *elpc-3* and *elpc-1* mutants on increasing amounts of mcm5S2U, of ncm5U or of both together (from 1 to 400 ⎧M).

Adding a single or the two modified Uridines to worm food had no effect on wt or *elpc* mutants chemo-attraction (not shown). To test if they could influence positive or negative imprinting, we odor-exposed wt and *elpc-1* worms in the presence of 1 to 400 ⎧M mcm5S2U, ncm5U, or both.

Wild-type imprinting was unchanged in the presence of any amount of one or both modified uridines (not shown). By contrast, we observed clear odor-specific effects of ncm5U in inhibiting the odor-triggered negative imprinting in *elpc-1* mutants (***Figure 13***). As already shown, (***Figure 10)***, odor-exposed *elpc-1* mutants acquire a stably inherited odor-specific inhibition of odor responses. The range of ncm5U required to inhibit *elpc-1* negative imprinting varies according to the odors. As low as 1 ⎧M of ncm5U impairs IA 300 imprinting, between 3 and 10 ⎧M impairs BA 300 imprinting, while between 30 and 100 ⎧M is needed to impair CI 300 imprinting. Upper or lower ncm5U concentrations, outside the respective odor-specific ranges, do not hinder negative imprinting. ***Figure 13*** data suggest that at least variations of the ncm5 uridine modification in tRNA^Ala^ (UGC) could participate to the formation of the odor-specific codes.

**Figure 13.**
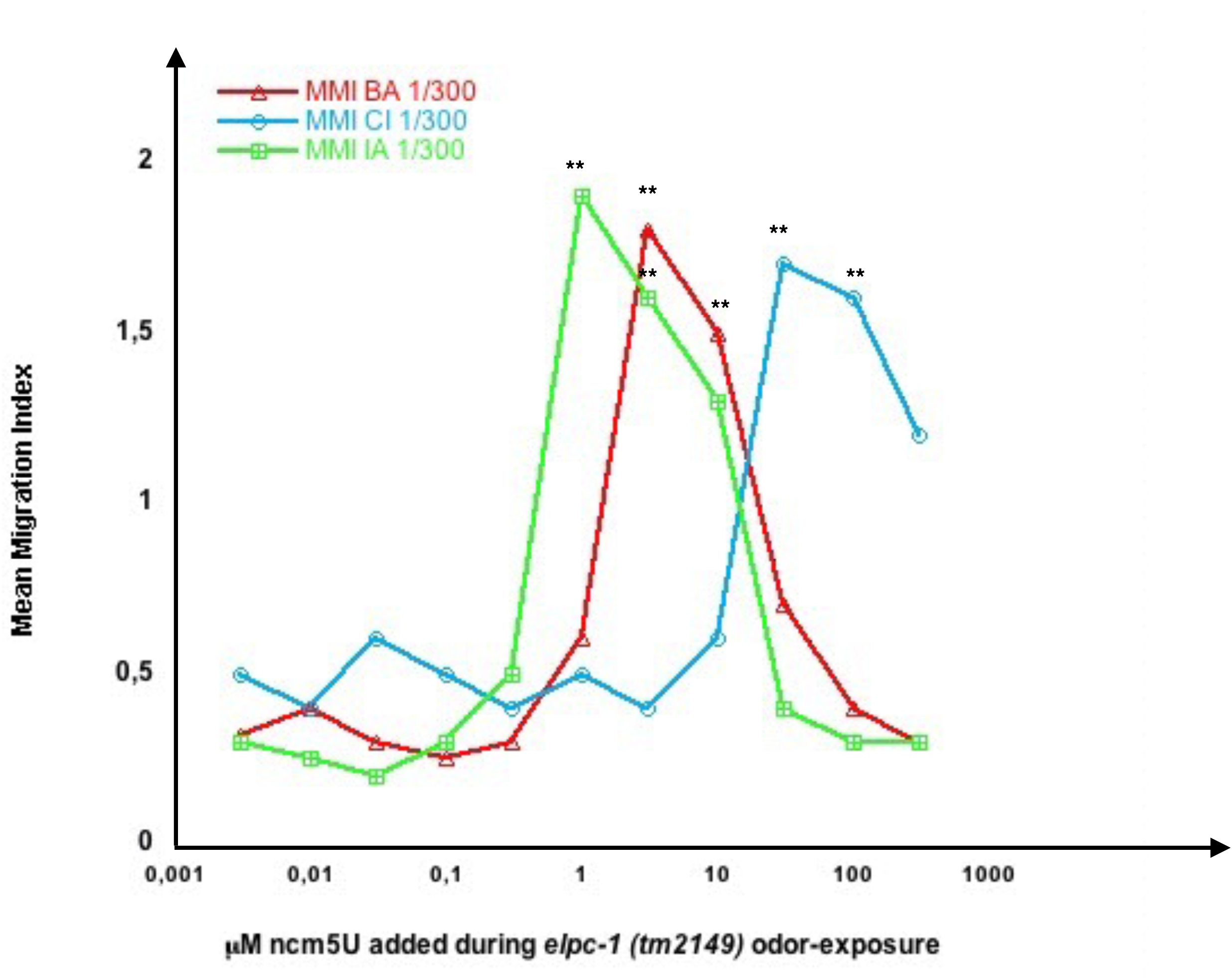
Odor-specific ranges of ncm5 modified Uridine impair odor-specific negative imprinting in *elpc-1* mutants. *elpc-1 (tm2149)* mutant worms were odor-exposed to either BA 1/300, CI 1/300 or IA 1/300 in the presence of increasing amounts of 5-carbamoylmethyluridine (ncm5U) from 0.004 to 400 μM. Mean migration Indices (MMI) in the respective BA, CI or IA odor gradients were determined at the adult stage, as described (experimental repeats ≥ 4, **p-value < 0.01).

In *C. elegans*, only two pairs of chemosensory neurons, AWA and AWC, are required for chemotaxis to volatile attractants (***Bargmann, 1993***). The *C. elegans* genome encodes around 1300 functional seven transmembrane receptors, presumably coupled to G-proteins, called Serpentine Receptors (SR). When expressed in chemosensory neurons, these receptors are thought to interact with odorant molecules and support odor specificity of the chemoattractive responses *(**Hart et al., 2010***). The AWC olfactory neurons are responsive to a high number of chemically different molecules that includes the three chemoattractants BA, CI and IA used in this study (***Bargmann, 1993***). Only the developing first larval stage L1 can be imprinted by attractive (this study), aversive (***Jin et al., 2016),*** or toxic stress ***(Gecse et al., 2019)*** signals present in worm’s environment. Environment-responsive mechanisms must be present at this stage to adapt future worm’s behaviors through up or down regulation of genes expressed in chemosensory neurons. A single chemo-attractive odor signal specifically increase, in wt, or decrease, in *elpc-1* mutants, adult attraction to a single odorant molecule, while all other AWC-mediated responses remain unchanged. Odor-specificity supports the existence of insulation of olfactory signaling within olfactory neurons, as already hypothesized for other forms of odor-induced adaptation in *C. elegans* (***O’Halloran et al., 2009; Juang et al., 2013***).

Naive unexposed *elpc-1* worms are odor-responsive, suggesting they express functional chemo-receptors. Although the outcome of single odor-exposure is to irreversibly decrease chemo-attraction, naive *elpc-1* mutants might still produce odor-specific tRNA^Ala^ (UGC) upon odor-stimulation. We hypothesized that the lowered odor-responses displayed by negatively imprinted *elpc-1* worms (***Figures 10 and 11***) could be due to lowered or null expression of the respective SR olfactory receptors. Without receptors, odors would not lead to the production of odor-modified tRNA^Ala^ (UGC). Naive *elpc-1 (ng10)* mutants are called CI ^+^ or BA^+^, while *elpc-1 (ng10)* that carry negative imprints of CI, BA or for CI and BA are respectively called CI ^-^, BA^-^ or CI ^-^/BA^-^. We fed N2 wt on tRNA^Ala^ (UGC) purified from naive, CI ^+^, CI ^-^, BA^+^, BA^-^and CI^-^/BA^-^ *elpc-1 (ng10)* worms, as indicated in ***Figure 14***.

**Figure 14.**
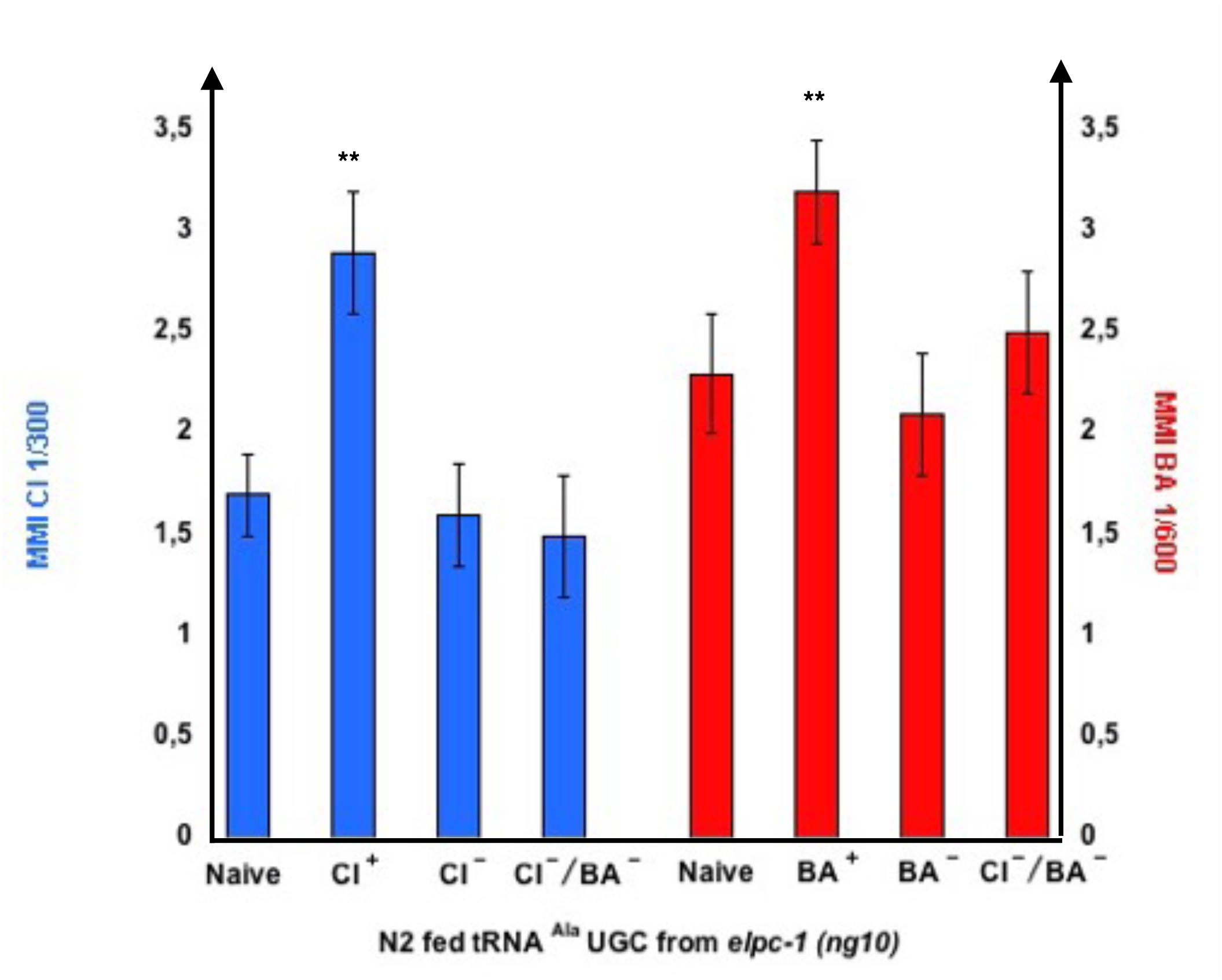
tRNA^Ala^ UGC from negatively imprinted *elpc-1 (ng10)* mutants do not transfer imprinting to naive. Stable reduction of CI response, BA response or CI and BA responses were obtained in *elpc-1 (ng10)* after odor-exposure or after odor-tRNA feeding, as described in Figure 11. Naive unexposed *elpc-1 (ng10)* worms are annoted C ^+^ and B ^+^ *elpc-1 (ng10)* worm populations with stable reduction of attraction to CI, to BA or to CI and BA are respectively annoted CI ^-^, BA ^-^ or CI ^-^/BA ^-^. N2 wt were fed on tRNA^Ala^ UGC respectively purified from *elpc-1 (ng10)* worms (left to right): Naive, CI-exposed, stable CI ^-^ CI-exposed, stable CI ^-^/BA ^-^ CI-exposed, Naive, BA-exposed, stable BA^-^ BA-exposed, and stable CI^-^/BA^-^ BA-exposed. MMI to CI 1/300 and to BA 1/300 were determined as described (experimental repeats ≥ 4,**p-value < 0.01).

As shown on ***Figure 14***, tRNA^Ala^ (UGC) from odor-exposed enhances chemo-attraction of wt worms, compared to tRNA^Ala^ (UGC) from naive unexposed. By contrast, tRNA^Ala^ (UGC) from worms bearing a CI ^-^ negative imprint or a BA^-^ negative imprint, do not, respectively, enhance CI or BA chemo-attractive responses of wt worms. As expected, *elpc-1 (ng10)* with a double CI ^-^/BA^-^ negative imprint do not enhance wt attraction to CI nor to BA.

The data from ***Figure 9b*** suggested *elpc-1* worms produce a wt form of tRNA^Ala^ (UGC), able to rescue the *elpc-3* chemotaxis phenotype.

From the results showed in ***Figure 14*** we can conclude that naive never-exposed *elpc-1* mutants produce the odor-specific tRNA^Ala^ (UGC) after odor-exposure, able to transfer a positive imprinting to wild-type. However, odor-stimuli do not anymore trigger the production of odor-specific tRNA^Ala^ (UGC) in negatively imprinted *elpc-1* worms. Whether or not negative imprinting is due to a down regulation of chemo-receptor expression remains however to be demonstrated.

Attractive odor stimuli activates an AWC-specific olfactory transduction pathway made of several G proteins, the DAF-11/ODR-1 guanylyl cyclase, and the TAX-2/TAX-4 cGMP-gated channel. Worms with mutations inactivating members of this pathway are defective for all AWC-mediated chemosensory responses, that include responses to the three attractive odorants used in this study (***Bargmann, 2006***).

We asked whether a functional AWC olfactory transduction pathway is required for the synthesis of odor-tRNAs. We purified tRNA^Ala^ (UGC) from CI-exposed worms harboring the *odr-1 (n1936)*, the *tax-2 (p671)* or the *tax-4 (p678)* mutations, respectively inactivating the ODR-1 guanylyl cyclase, the TAX-2 or the TAX-4 subunits of the cGMP-gated channel. Wild-type and *elpc-1 (ng10)* mutants were then fed on these tRNAs, in order to compare their imprinting activities ***(Figure 15)***.

**Figure 15.**
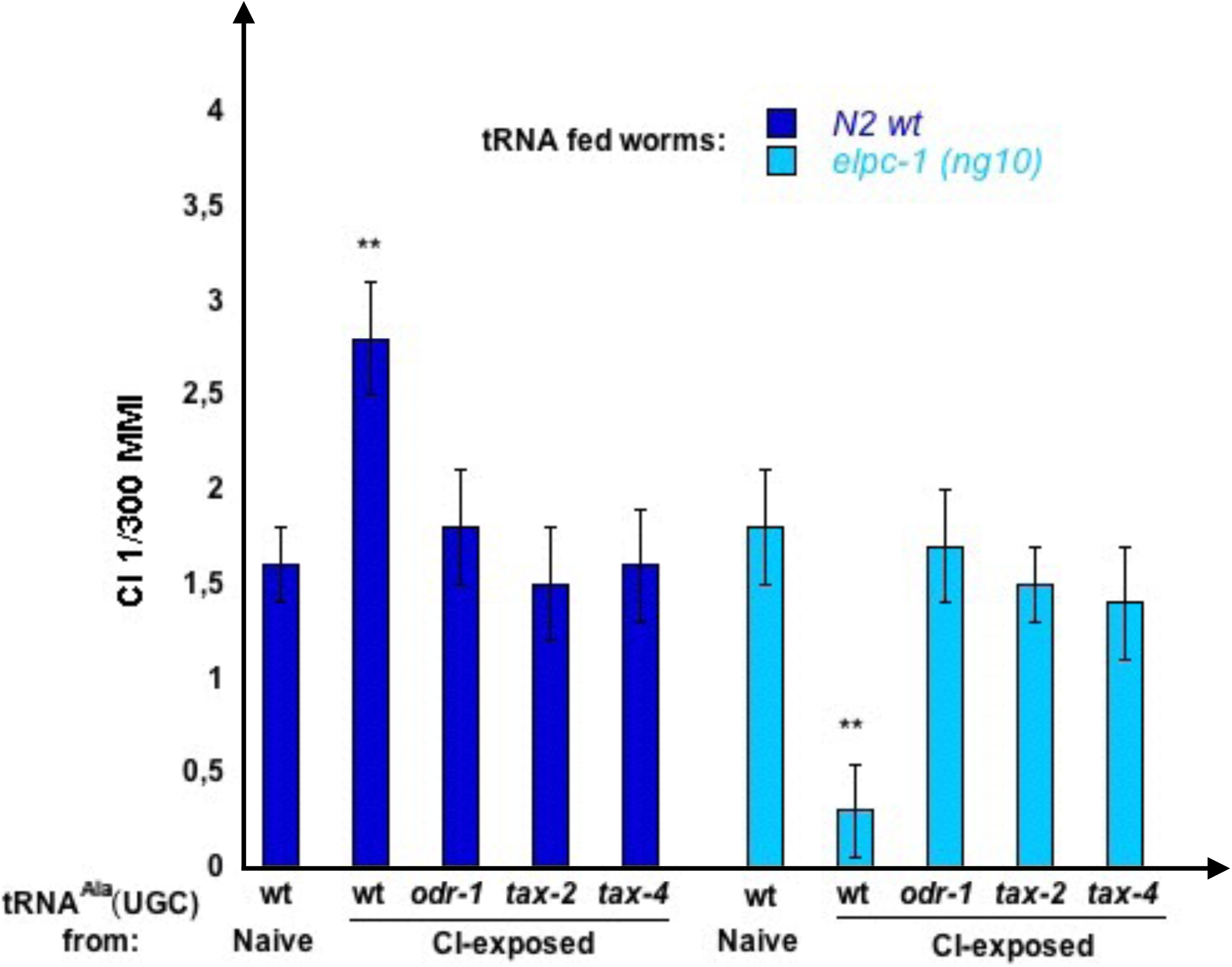
Olfactory transduction mutants do not produce odor-imprinting tRNA^Ala^ (UGC) after odor-exposure. tRNA^Ala^ (UGC) was microbeads purified from Naive unexposed, CI 1/300-exposed wt N2 worms, CI 1/300-exposed *odr-1 (n1936),* CI 1/300-exposed *tax-2 (p671)*, and from CI 1/300-exposed *tax-4 (p678)* mutant worms. Naive wt and naive *elpc-1 (ng10)* worms were fed on these tRNA^Ala^ (UGC). Mean Migration Index to CI 1/300 was determined at the adult stage as described (**p-value < 0.01).

As already shown, feeding on tRNA^Ala^ (UGC) from CI-exposed wt increase CI MMI in wt (positive imprinting), and decrease CI MMI in *elpc-1 (ng10)* (negative imprinting). By contrast, none of the three CI-exposed olfactory transduction mutants produce a tRNA^Ala^ (UGC) able to transfer a positive CI-imprint to naive wt or a negative imprint to *elpc-1 (ng10) (*CI-exposed *odr-1*, *tax-2, and tax-4)*.

Together, ***Figures 14 and 15*** results indicate that imprinting requires chemo-receptors to be expressed and coupled to a functional odor-transduction pathway, as early as the imprinting L1 critical stage. They also support the idea that the production of odor-specific tRNA^Ala^ (UGC) occurs downtream of the odor-signaling.

## DISCUSSION

Our data indicate both the innate and the experience-modulated chemo-attractive behaviors are under the control of a single tRNA molecule. To our knowledge, this is the first report demonstrating a behavior is under the control of a transfer RNA.

The tRNA^Ala^ (UGC) would carry specific chemical codes required for the acquisition of chemo-attractive responses, and for the transmission of odor-specific information. As other RNAs, tRNAs can adopt many chemical forms by combining more than 160 chemically modified bases (***Boccaletto P. et al., 2018)***. Combinatorial calculation indicates that even if just a fraction of the nucleotides are chemically modified, the number of possible combinations remains very high. Theoritically, choosing 10 base modifications (m) out of 100 (n) already offers more than 10^13^ possible combinations, according to the n + m - 1)! / ((n - 1)! * m! equation. To understand how such huge potential of chemical diversity is translated into functional diversity represents a considerable challenge.

Although mature tRNAs mainly control protein translation, tRNAs can be fragmented into smaller RNA molecules. tRNA fragments, tRF, are an emerging class of non-coding RNAs that may, besides translation, control several other important biological functions (***Megel et al., 2015; Pliatsika et al.; 2018***).

The 3D structure and bases modifications contribute to the high stability of mature tRNAs (***Engelke and Hopper, 2006***). It would be interesting however to analyse if and how mature tRNAs could be processed after contact with the bacterial lawn (worm food), and after ingestion and diffusion from intestinal cells to other tissues, including germ-line cells. It has been already shown that endogenous RNAi in the chemosensory neuron AWC promotes odor-specific adaptation in adult *C. elegans* (***Juang at al., 2014***). If tRF can be generated by Dicer-independent mechanisms, the tRNA double-stranded stretches could be processed via well described Dicer-dependent small non-coding RNA pathways. If it is the case, genetic inactivation of these pathways would impair the tRNA^Ala^ (UGC) triggered biological activities described in this paper. Moreover, comparative RNA sequencing after tRNA^Ala^ (UGC) feeding may help to identify which small RNAs eventually originated from tRNA processing could trigger the epigenetic changes.

Each nucleotide chemical modification requires the activity of specific enzymes, called writers. So far, no *C. elegans* tRNA writers are known, except TUT-1 and the Elongator complex, both modifying Uridines, respectively, into S2U, mcm5S2U and ncm5U. Through the addition of different amounts of these three modified Uridines to worm food during odor-exposure, we identified ncm5U as being part of the odor-specific codes carried by tRNA^Ala^ (UGC). Providing worms with different amounts of ncm5U however only impairs, thus odor-specifically, the negative olfactory imprinting in *elpc-1* mutants, but does not affect the positive imprinting in wild-type. We hypothesize that specific patterns of base modifications would stand for specific biological activities. Transfer and inheritance may rely on yet unknown reading mechanisms, able to recognize the modification patterns, and of copying mechanisms, able to precisely reproduce those.

We showed that adding the wild-type tRNA^Ala^ (UGC) to the food of *elpc-3* mutants rescues their chemo-attractive defects. Rescue could rely on compensation of the lacking modified U by those carried by the wt tRNA.

However, adding the « defective » *elpc-3* tRNA^Ala^ (UGC) to wt worms phenocopy the *elpc-3* behavioral phenotype. In this case, compensation of lacking modifications cannot account for the transferred phenotype, suggesting phenocopy could be rather based on recognition and reproduction of the « defective » modification pattern in the mutants. The mechanisms by which epitranscriptomic codes are written, red, copied or erased, remain elusive. Analysing the activity of a tRNA^Ala^ (UGC) made by wild-type worms fed on tRNA^Ala^ (UGC) made by known modification mutants, as *tuc-1*, and inversely, could provide some further insights.

We report that deletions in the *elpc-3* gene do not have the same behavioral effects as deletions of the *elpc-1* gene. Our observations suggest that the catalytic sub-unit ELPC-3 modifies tRNAs in the absence of a functional ELPC-1, while ELPC-1 is required for imprinting. They challenge previous reports in which inactivation of any elongator sub-unit produced the same phenotypes. However, the structural organization and the respective biochemical functions of the *C. elegans* Elongator sub-units are unknown. Whether the multiple elongator functions only rely on its tRNA modifying activity, or on other biochemical activities of its sub-units, is still debated. As indicated in WormBase(***<u>wormbase.org</u>***), the comprehensive resource for nematode research, *elpc-1* genetically interacts with *chaf-2* and *dhc-1*. CHAF-2 is an ortholog of the human chromatin assembly factor CHAF1B, involved in nucleosome assembly, linking newly synthesized histones deposition to DNA replication **(*Sauer et al., 2018*)**. DHC-1, the unique *C. elegans* cytoplasmic dynein 1 HC, controls microtubule dynamics, and is involved in many essential biological processes, including neuron development and meiotic spindle orientation (***O’Rourke et al. 2007***). These two interactions could be particularly relevant to the *elpc-1* mutant phenotype described here, which associates tRNA metabolism, epigenetic reprogramming and neuro-development.

Nematodes respond to many volatil odorants but has only three pairs of olfactory neurons, AWA, AWB and AWC, suggesting each neuron would express a large number of odor-sensitive chemo-receptors. It has been shown that the type of behavioral response elicited by an odorant is not specified by the chemo-sensory receptors, but by the olfactory neuron in which they are expressed and activated **(*Troemel et al., 1997*)**. Our results suggest chemo-receptors for attractive cues are expressed and already functional, i.e. coupled to the olfactory transduction pathway, at the first stage of larval development. The way each worm’s chemo-responsive neuron expresses specific sets of receptors, and represses the expression of all others, is still unknown. In mammalian species, each olfactory neuron expresses only one allele of a member of the olfactory receptor genes family; the precise mechanisms by which such monogenic monoallelic receptor choice is achieved is however not fully understood. These mechanisms include nuclear compartimentalization, repression/derepression through epigenetic marks as those made by the chromatin-modifying enzyme Lysine demethylase 1 (LSD-1) and stabilization of functionally expressed receptors by the odor-signaling pathway (reviewed in ***Degl’Innocenti A and D’Errico A, 2017***).

tRNAs and tRNAs genes not only control translational ***Kirchner and Ignatova, 2015; Avihu et al., 2013*** but also transcriptional fine-tuning (Woolnough et al., ***2015; Pratt-Hyatt et al., 2009; Kirkland and Kamakaka, 2015***. Importantly for epigenetic inheritance, tRNAs also control the plasticity of genome architecture via the dynamic positioning of nucleosomes and the modulation of chromatin domains boundaries ***Raab et al., 2012; McFarlane and Whitehall, 2009; Talbert and Henikoff, 2009***).

In this work, we produced worm strains with stably inherited non-genetic alterations of odor-specific responses, compared to wild-type. Increased or decreased odor-specific attraction may rely on the differential expression of odor-specific receptors. Through combinatorial epigenetic modifications, the nuclear localization of their genes may evolve, moving from either active or repressed chromatin domains (***Armelin-Correa et al., 2014; Evans et al., 2016***).

The pattern of chemo-receptors expression in *C. elegans* sensory neurons is unstable as it can be altered by life history and external conditions (***Vidal et al., 2018***). Worms are innately attracted by a number of volatil molecules, however only within a limited range of dilutions. The mechanisms that specifically direct attraction to each odor-specific dilution might combine sensory neuron distribution and differential expression levels of specific sets of chemo-receptors.

Knowing which genes are involved in an odor-specific response is a prerequisite to further delineate the epigenetic mechanisms that would regulate expression patterns and levels. The stably odor-specifically imprinted worms described here, displaying permanent higher or lower responses to single or to multiple odorants, compared to wild-type naive, might greatly facilitate the identification of these genes.

This work highlights the role of a specific tRNA and the multifunctional Elongator complex at the interface between environmental inputs and behavioral changes. It sets the stage to elucidate how environmental informations, here olfactory cues, could regulate secondary and tertiary tRNA structures, which, in turn, will influence worm behavior.

Due to the growing number of human diseases linked to RNA modification defects (***Sarin and Leidel, 2014***), understanding the biological significance of the epitranscriptome is a major issue. Recent findings strongly indicate that RNA modifications play dynamic regulatory roles that are analogous to the epigenetic modifications of DNA and histone proteins. Understanding the function and mechanisms of the dynamic RNA modifications, or epitranscriptomic, represents a new challenge at the frontier between different disciplines. Reversible RNA modifications add a new dimension to the developing picture of post-transcriptional regulation of gene expression. This new dimension awaits integration with transcriptional regulation to decipher the multi-layered information that controls a plethora of biological functions. Understanding how non-coding transcriptomes are modified in response to environmental changes and how these modifications impact germ-line cells and translate into inherited phenotypes thus represents an important challenge (***Schimmel, 2018***).

Olfactory imprinting serves hatchling attachment and adult homing to the native chemo-sensory environment in many animal species (***Grassman et al., 1984; Dittman and Quinn, 1996; Horn, 2004; Gerlach et al., 2007; Gerlach et al., 2019***).

As we describe here for the nematode, multigenerational imprinting of the same olfactory cues would lead to stable inheritance. Through a process of (epi)genetic assimilation, different animal populations, though belonging to the same species, may have acquired specific patterns of chemo-attractive responses, only depending on their local living environments. This kind of « cultural » differenciation, through the stable assimilation of responses to external challenges, may be instrumental to the evolution of innately expressed behaviors (***Jablonka and Lamb, 2015)***.

Our data is consistent with the model presented in ***Figure 16***. In *C. elegans,* the first post-hatching hours are highly receptive to environmental conditions (***Hall et al., 2010; Jin et al., 2016; Hong et al., 2017; Gesce et al., 2019***). Imprinting chemo-attractive cues is an innate behavior taking place during this period of development. Its biological function would be to confirm that olfactory cues present in the early environment are indeed encoded as innately attractive and will be potentially rewarding for the future worm life and for its progeny.

**Figure 16.**
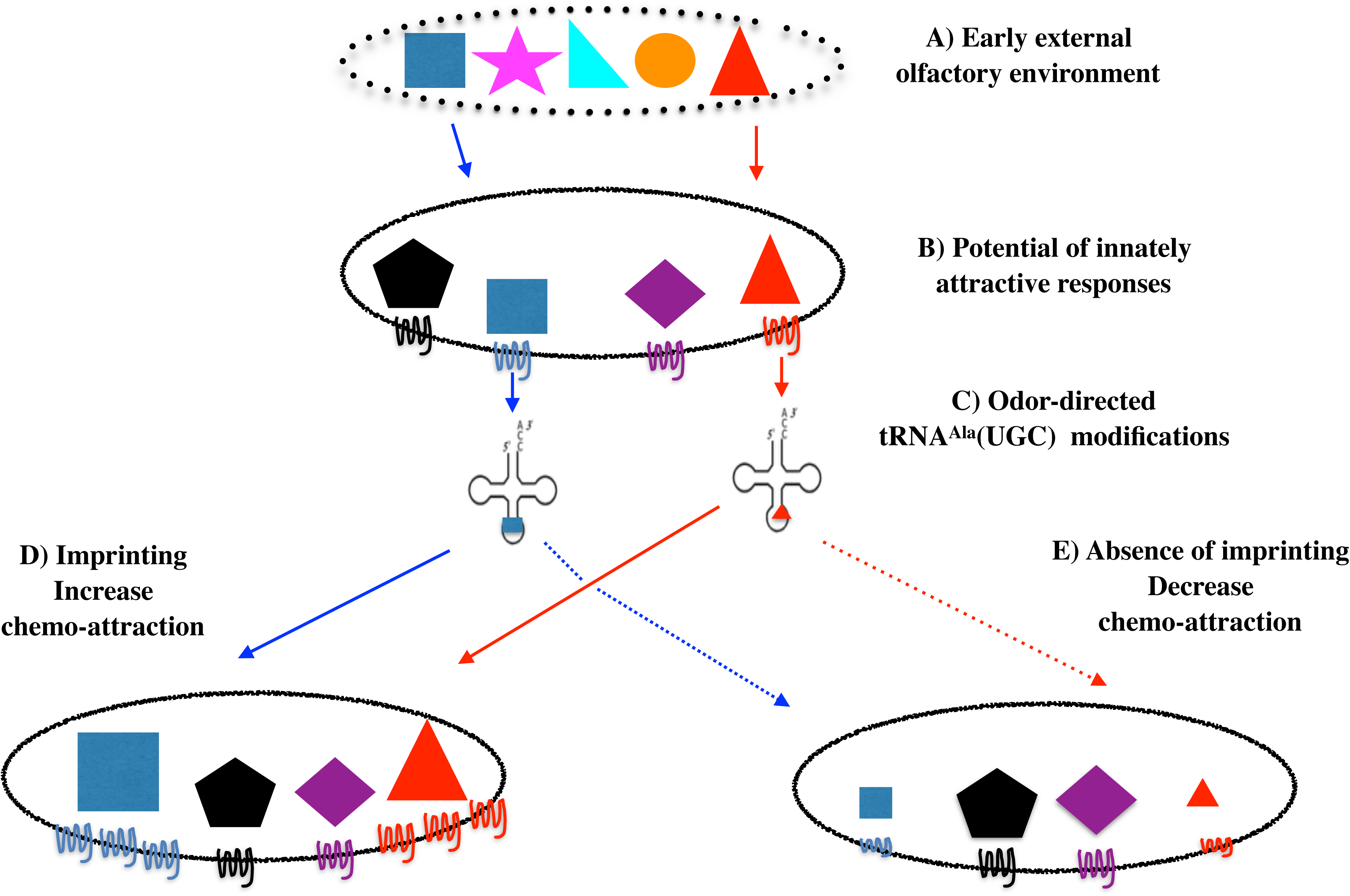
Imprinting is an innate behavior that sanction innately encoded chemo-attraction and early environment adequacy. The olfactory environments in which animals come to life contains a number of chemicals (A). Worms are innately attracted by some of these chemicals, here represented by the blue square and the red triangle (B). (C) Attractive odors trigger the production of tRNA^Ala^ (UGC) carrying odor-specific information. (D) Imprinting will increase worm attraction to odorants that are both present in the early environment and encoded as innately attractive. (E) In the absence of imprinting, as in *elpc-1* mutants, odorants will stably decrease the innate worm responses for these odorants.

An innate frame would encode and program the whole potential of *C. elegans* chemo-attractive responses. Qualitative (to which chemicals) and quantitative (attractive strength) responses are stably inherited over generations in stable environments. As environment changes, it may or not contain cues encoded as innately attractive. If it does not, the spectrum of behavioral responses will not change as it relies on the innate program.

If it does, external olfactory cues will interact with the chemo-receptive elements set up to transduce chemo-attraction. Such interactions would trigger odor-specific labelling of tRNA^Ala^UGC, using the combinatorial potential of tRNA bases modifications. In this diffusible form, odor-specific information can move internally, from neuronal to, eventually, germ-line cells, and, as we show here, transfer olfactory imprints from experienced to naive animal. Such diffusible mnemonics would be a way to encode, transfer and communicate experience without modifying the neuronal synaptic network. Moreover, they could support the transfer of experience in biological organisms with low number or without neurons (***Boussard et al., 2019***). Reading mechanism should exist that translate the epitranscriptomic codes into unstable or stably inherited epigenetic codes, including histone modification codes, leading to transient or stable gene expression and behavioral changes. By enhancing their attractiveness, the role of imprinting would be to confirm that the newly encountered cues are well assessed.

However, even in the case environmental signals and innate program coincide, attraction can not be confirmed in imprinting defective animals, as we showed here for *elpc-1* deleted mutants; without confirmation by the imprinting process, worms would be definitely desensitized and loose attraction to the specific cues they have been exposed to. Imprinting is thus an innate behavior that supports plasticity of the innate directory of behaviors.

## Acknowledgements

We thank Craig Hunter for the *sid-2 (qt13)* mutant. The elongator mutants *elpc-1 (ng10)* or *ikap-1 (ng10)* and *elpc-3 (tm3120)* were obtained thanks to Jachen A. Solinger via Grazia Malabarba. *elpc-3 (tm3120)*, *elpc-1 (tm2149)* and *elpc-1 (tm11684)* knockouts were generated by the National Bioresource Project, Tokyo, Japan, which is part of the International C. elegans Gene Knockout Consortium. We thank Sebastian Leidel for providing the *tuc-1 (tm1297)* worms. Other strains used in this study were provided by the Caenorhabditis Genetics Center, which is funded by the National Institutes of Health (NIH) Office of Research Infrastructure Programs (P40 OD010440). We thank Wormbase for providing valuable databases. Thanks to V. Grandjean and G. Raad for stimulating discussions and incentive suggestions. This work was supported by a grant of the “Agence Nationale de la Recherche” ANR-12-Bioadapt-0022, and by the LABEX ANR-11-LABX-0057_MITOCROSS program.

## Material and Methods Modified Uridines

S2 Uridine (S2U) was obtained from Jena Bioscience (Germany) and diluted in water. 5-methoxycarbonylmethyl 2-thiouridine (mcm5S2U) and 5-carbamoylmethyluridine (ncm5U) were custom synthesised by Jena Bioscience, and diluted from 20 mM stocks in Tris 0.3 M pH 8.

### Strains and culture conditions

We used the wild-type *C. elegans* reference strain N2, unless otherwise stated. Worms with the *sid-1 (qt2)* and the *elpc-3 (ok2452)* mutations were obtained from the *Caenorhabditis* Genetics Center (CGC). The *sid-2 (qt13)* mutant was obtained from Dr C. Hunter. The 375bp deletion in the *elpc-3* gene called here delKAT was produced by Nemametrix, Inc. (Eugene, OR, USA) using the CRISP-sdm transgenic method. Deletion was confirmed by PCR/RE and sequencing. The elongator mutants *elpc-1 (ng10)* (or *ikap-1 (ng10)),* and *elpc-3 (tm3120)* were from Dr J. A. Solinger, while the *elpc-1 (tm2149)* and *elpc-1 (tm11684)* deletions were from Dr S. Mitani/NBRP. The *tuc-1 (tm1297)* mutant was from Dr S. Leidel. Worms were cultured on E. coli OP50 at 20°C using standard protocols.

### Odor-exposure for olfactory imprinting

Benzaldehyde, -citronellol or Isoamyl alcohol (Sigma-Aldrich) were diluted as® described in water. Odor-exposures were done by suspending a 4 µl drop of these dilutions on the lids of worm culture dishes at least during 24 hours from the egg stage at 20°C, covering the critical plasticity period corresponding to the first 12 hours of post-hatch development (***Remy and Hobert, 2005***).

### Chemotaxis assay

The chemotaxis assay used in this study is schematically outlined in the Supplement Figure Method. It is based on the population chemotaxis assay originally described by C. I. Bargmann (***Bargmann, 2006***). Several modifications were made in the procedure and in chemotaxis index calculation. Changes aimed at more accurately compare the chemo-attraction of worm populations to moderately attractive odorants (as the dilutions of attractive odorants used in this study), using 20 adult worms per condition.

40 ml of low-salt (1mM Ca^++^, 1mM Mg^++^, 5mM KPO_4_) agar (20 g/l) were poured in 12×12 cm square plates. Assay plates were allowed to dry at room temperature for at least three days before use. Worms were individually transferred from culture dishes to the middle of the square assay plates. To establish a homogeneous odor gradient, so that all worms were submitted to the same olfactory stimulus, 3 drops of odor-dilution (4 µl each) were suspended on the lids at one side, each placed at a distance of 3 cm from the others. At the opposite side of the lids were placed 3x 4 µl of water. 6 drops of 4 µl of 1M NaN3 were added at both sides of the agar plate to immobilize animals that reached edges.

At time 0, 20 worms were placed on the middle line of the squared plate, every worm being at a distance of 6 cm from odor sources. Assays performed on squared plates allowed the indexation of all worm positions between the starting line (time 0, position 0 cm) and the odor source (position + 6 cm), usually four times at 10, 20, 30 and 40 minutes from time 0. The mean value of all indexed positions (in cm from the starting line) of each of the 20 worms represented the Mean Migration Index (MMI). Each experiment shown in the paper was performed at least 4 times. MMI (Means +/-S.E.) values were compared using unpaired data with unequal variance Student t-tests performed with the KaleidaGraph program. Assays were always performed so as to compare synchronized worm populations.

### RNAs fractionation and tRNA purification

The large and small RNAs (< 200 nt) were separated using the NORGEN RNA Purification Kit (Norgen Biotek Corp). Small RNAs were further size-fractionated on 3.5% low-melting agarose (Nusieve GTG) gels. RNA was quantified with the Nanodrop 2000 (ThermoScientific). To purify *C. elegans* transfer RNAs, small RNA fractions (under 150 nt in size) were prepared as described (***Maréchal-Drouard et al., 1995***). In brief, RNA was incubated in a 2 M lithium chloride solution over-night at 4°C. After a centrifugation at 16000 g for 30 min at 4°C, the supernatants containing the tRNAs were recovered. Transfer RNAs were thereafter precipitated with 0.1 volume of 1 M sodium acetate pH 4.5 solution and 2.5 volumes of ethanol over-night at −20°C. Pelleted tRNAs were dissolved in water and loaded on 15 % polyacrylamide with 7 M Urea and 1 X TBE gels. After Ethidium bromide staining, gel slices were cut from the gels. The tRNAs were eluted over-night at room temperature in a solution composed of 0.5 M ammonium acetate, 10 mM magnesium acetate, 0.1 mM EDTA and 0.1% SDS. After phenol extraction, tRNAs were ethanol precipitated and finally recovered in 10l water. ⎧

### Microbeads-coupled tDNA probes for tRNA isolation

To purify tRNA molecules, we used the microMACS Streptavidin MicroBeads Kit (Miltenyi Biotec). We synthesized 37 nt long 3’-biotinylated DNA probes complementary to the 3’ half of the *C. elegans* tRNAs. Oligonucleotide sequences of the tDNA probes were deduced from the *C. elegans* tRNA genes predictions (GtRNAdb, Lowe lab, Biomolecular Engineering, University of California Santa Cruz). The sources of RNA from which specific tRNAs were isolated were either the whole small RNA populations or the gel-eluted tRNA fractions showed in ***Figure 3***. Procedure was as recommended by the microMACS Kit, except the annealing buffer was made of 10 mM Tris HCl pH 7.5, 5 mM MgCl_2_. Binding/Wash buffer 5 X was 50 mM Tris HCl pH 7.5, 5 mM EDTA pH 8, and 2.5 M NaCl, as indicated. Elutions from microbeads were in 200 l TE. For each feeding and behavioral assay, we used ⎧ 10 eluate per worm culture dish, as described in the RNA-feeding section below. ⎧l

### High purification of tRNA molecules after **a**ffinity chromatography and 3’ pCp-Cy3 labeling

To obtain highly purified Alanine tRNA (UGC) molecules, we used the Streptavidin Sepharose^TM^ High Performance beads (GE Healthcare, 17-5113-01) coupled to the 3’-biotinylated tDNA Ala (TGC)-3’ probe (probe Ala TGC N° 3, as described).

We first mixed 10-20 µg of total tRNA with 1 nmole of tDNA probe and 40 U of RNaseOUT in 500 µl of annealing buffer (5X SSC (saline-sodium citrate); 0.05% SDS (sodium dodecyl sulfate)). The mix was then incubated sequentially, 5 min at 70°C, 30 sec at 4°C and 2.5 hours at 42°C. Twenty µl of Streptavidin Sepharose beads were deposited in a 20 µm receiver column (Macherey-Nagel, 740522) and equilibrated with 3x 500 µl of annealing buffer. Columns were incubated 15 min at room temperature in gentle shaking and 15 min at 42°C. They were centrifuged 15 sec at 11000 *g* and then washed as follows: 2x 500 µl of 5x SSC (5 min at 42°C), 1x 500 µl 2x SSC (5 min at 42°C), 1x 500 µl 2x SSC (15 min at 47°C), 1x 500 µl 2x SSC (15 min at 52°C). After each washing step, the column was centrifuged 15 sec at 11000 *g*. tRNA elution was by adding 2x 300 µl of elution buffer (10 mM Tris-HCl pH 7.5; 1 mM EDTA; 5 M Urea). Total elution was ethanol precipitated and resuspended in 2.96 µl of water for 3’ pCp-Cy3 (Jena Bioscience NU-1706-CY3) labeling.

Before labeling, we added 26 % (v/v) DMSO (1,04 µl) to the purified tRNA (2,96 µl) and incubated 10 sec at 100°C and immediately cooled on ice. The labeling reaction was performed using the Biolabs T4 RNA ligase (M0204). Ligase buffer and ATP were added to the tRNA-DMSO mix and ligation was performed with 20 µM of pCp-Cy3 and 5 units of T4 RNA ligase in 10 µl at 16°C overnight. For total tRNAs fraction, the labelling was done in the same way, using 2.5 µg total tRNAs.

The different fractions were analyzed by electrophoresis on 7 M Urea-15 % acrylamide gel and scanned on a GE Healthcare Ettan DIGE imager system. The bands corresponding to the purified tRNA were cut from the gel and eluted overnight at room temperature in 0.5 M ammonium acetate, 10 mM magnesium acetate, 0.1 mM EDTA and 0.1% SDS. After phenol extraction, tRNAs were ethanol precipitated and finally recovered in 8 µl of water.

### Northern blots analyses

Northern blots were performed as described in ***Cognat et al., 2017***.

Roughly, RNAs fractionated on a denaturing 7M Urea/15 % polyacrylamide gel were transferred onto Hybond-N+ membrane (Amersham Pharmacia Biotech). Membranes were then hybridized with **^32^**P labeled probes specific to *C. elegans* tRNAs:

tRNA**^Ala^**(AGC): 5’- CTACCACTGAGTTATACCCCC - 3’
tRNA**^Ala^**(CGC): 5’- TACCCCTGAGCTATACCCCC - 3’
tRNA**^Ala^**(UGC): 5’- TATGGGGAATCGAACCCCA - 3’
tRNA**^Leu^**(AAG): 5’- TGGTGAAGAGAGTGGGATTCGAAC - 3’
tRNA**^Gly^**(UCC): 5’- TGGTGCGTTCGGGGGGAATCGAAC - 3’
tRNA**^Lys^**(UUU): 5’- ACCAACTGAGCTAAGGAGGC - 3’
tRNA**^Glu^**(UUC): 5’- AACCACTAGACCACATGGGA - 3’
tRNA**^Gln^**(UUG): 5’- AACCGCTACACCATGGAACC - 3’

### RNA-feeding

NGM agar plates were loaded with 40 µl of OP50 culture, which, after drying, formed approximately a 100 mm^2^ spot. Volumes of 10 µl RNA to be assayed were deposited per *E. coli* spot and shortly dried. Synchronised naive embryos were spawned on the RNA-loaded plates, and worms grown at 20°C until adulthood. For imprinting inheritance after multi-generational tRNA feeding ***(Figure 5)***, N2 worms were grown from the spawn embryo to the adult laying stage on 1 ng CI-tRNA-loaded cultures plates (F1). Part of the next generation was grown on a new CI-tRNA-loaded culture plates - the second tRNA-fed generation F2 -, another on a regular plate without CI-tRNA - the first naive generation F1+1. F3 is the progeny of F2 grown on tRNA while F2+1 is the progeny of F2 grown without tRNA. The CI chemo-response of the naive generations from the seven successive tRNA fed generations (F1 to F7) was determined as described.

### tDNA oligonucleotides probes used for microbeads tRNAs purification

1. Ala AGC: 5’ TGGAGGTTTGGGGAATTGAACCCCAGCCCTCTCCCAT 3’
2. Ala CGC: 5’ TGGAGGCACGGGGGATTGAACCCCGGACTTCCCGCAT 3’
3. Ala TGC: 5’ TGGAGGTATGGGGAATCGAACCCCAGACTTCTCGCAT 3’
4. Ala TGC: 5’ ATGCAAAGCCAGCGCTCTACCCCTGAGCTATACCCCC 3’
5. Arg ACG: 5’ CGACCACGGCAGGATTCGAACCTACAATCTTCTGC 3’
6. Arg CCG: 5’ AGCTCGCGGAGGGACTTGAACCCCCATTCCCGGTTCC 3’
7. Arg CCT: 5’ CGACCGAGGCAGGACTCGAACCTGCTGTCTTCGGTTT 3’
8. Asn GTT: 5’ CGCTCCCGGTGGGCTCGAATCACCTTTCGGTTAA 3’
9. Asp GTC: 5’ CTCCCCGGCCGGGAATTGAACCCGGGTCTCGCATGTG 3’
10. Cys GCA: 5’ CTAGCTCTCCAGGGACCAAGTTGAGGCCCACGGGGGA 3’
11. Gln CTG: 5’ CTTAGGACGCTGGGCTCAAGTTTAGAGCCACCCTGGA 3’
12. Gln TTG: 5’ CTTAGGACGCTGGGCTCAAGTTTAGAGCCACCTTGGA 3’
13. Glu CTC: 5’ GTGGGTATTCCGGCCCCAAGCTAAGGGGCGTTGCTTT 3’
14. Glu TTC: 5’ GTGGGTGCGCCGGGCCCAAGCTAAGGGCCGTACCCTT 3’
15. Gly CCC: 5’ ACCATTGTCTCGCGCCCAAGCTTAGGGCAGGTGGCGT 3’
16. Gly TCC: 5’ GTTCGTAAGCTGCCCCCAAGCTAAGGGGAGCTTGCGT 3’
17. His GTG: 5’ CCGGCACCGCTGCGACCAAGCTAAGGTCGTCGTCCGT 3’
18. Ile AAT: 5’ TTCGGGTTCCAGCGTCCAAGCTGGGGACGACCGCCGT 3’
19. Ile TAT: 5’ CAATTTGGTCAGCGCCCAAGCTTAGGGCGGGCCCCGT 3’
20. Leu AAG: 5’ GGGAAGCCCCCGCACCCAAGCTTAGGGAGAGAGAAGT 3’
21. Leu CAA: 5’AGCATACCCACGCACCCAAGCTTAGGGTGAAGCACGT 3’
22. Leu CAG: 5’AGAGGCCTCCCGCGTCCAAGCTTAGGACGCCTGCCGT 3’
23. Leu TAA: 5’ GGGAGGCCCCCGCACCCAAGCTTAGGGTGAGAGTAGT 3’
24. Lys CTT: 5’ ATTAGACCAACAGCGCCAAAGCTCGGGGCGTAACCCA 3’
25. Lys TTT: 5’ TTAGAATTCCAGTCCCCAAGCTCAGGGGATCCACCGA 3’
26. Met CAT: 5’ TTGGGTCTCCAGCCACCTAGCTTTGGTGAGCGACGAT 3’
27. Met CAT: 5’ TTAGACTTCCAGCACTCAAGCTCGGAGTGGCCCTCGT 3’
28. Phe GAA: 5’ TTTAGCAATCCAGTGGTCAAGCTAGGACCAAGCCCGT 3’
29. Pro AGG: 5’ CACGTTCTCTAGGGCCCAAGCTAGGGGCCAAGCTGGG 3’
30. Pro CGG: 5’ CACGCTCTCCAGGGCCCAAGCTAAGGGCCAAGCCGGG 3’
31. Pro TGG: 5’ CACGCTCTCCAGGGACCAAGTTAGGGGCCAAGCCGGG 3’
32. Ser AGA: 5’ CCGAGACGGGCGCATCCAAGCTTAGGACGACTGACGC 3’
33. Ser CGA: 5’ CCGAGACGGGCGCATCCAAGCTTAGGACGACTGACGC 3’
34. Ser GCT: 5’ CCCAAAGGGGCGCACTCAAGCTTAGAGTAGAACTAGC 3’
35. Thr AGT: 5’ TTTGTCTTCCAGCGACCAAGCTAAGGTCGTACTCCGT 3’
36. Thr CGT: 5’ TTTGTCTTCCAGCTGCCAAGTTAGGGCAGACCCCCGT 3’
37. Thr TGT: 5’ TAGTTATCCAGGCCCCAAGCTGGGGAGCATTCCCAGT 3’
38. Trp CCA: 5’ CTAGCTTTCCATCCCGCAAGCTAGGCGAGTCACCAGT 3’
39. Tyr GTA: 5’ TAGGAATCCAGTGACCAAGCTTAGGCCAAGCTGCCT 3’
40. Val AAC: 5’ TGTGTCTTCCAGCCACCAAGCTCGGGCGGGCTCTAGT 3’
41. Val CAC: 5’ TGCGTCTTCCAGCGGCCAAGCTTGGGCCGGTCCTGGA 3’
42. Val TAC: 5’ TGCGTCTTCTAGCGGCCAAGCTTGGGCCGATCCTGGA 3’

## Supplement Figure Method 1

### Schematic outline of the chemotaxis assay

Chemotaxis assays were performed on square plates in order to index the worms’ positions between the starting line (time 0, position 0 cm) and the odor source (position + 6 cm) or water source (position - 6 cm). 3 ml of odor solution or water are added to the lids of the plates. 20 synchronised worms are placed at the starting line at time 0. Worms start their migration and their position is carefully noted, usually at four time-points, at 10, 20, 30 and 40 minutes past time 0. Worms that migrate up the odor gradients are positively indexed (from 0 to 6). Worms that migrate down the odor gradients towards water are negatively indexed (from 0 to – 6). The mean value of all indexed positions at all noted times (in cm from the starting line) of each of the 20 worms involved in the assay is the Mean Migration Index (MMI).

---

**Supplement Figure 1.**
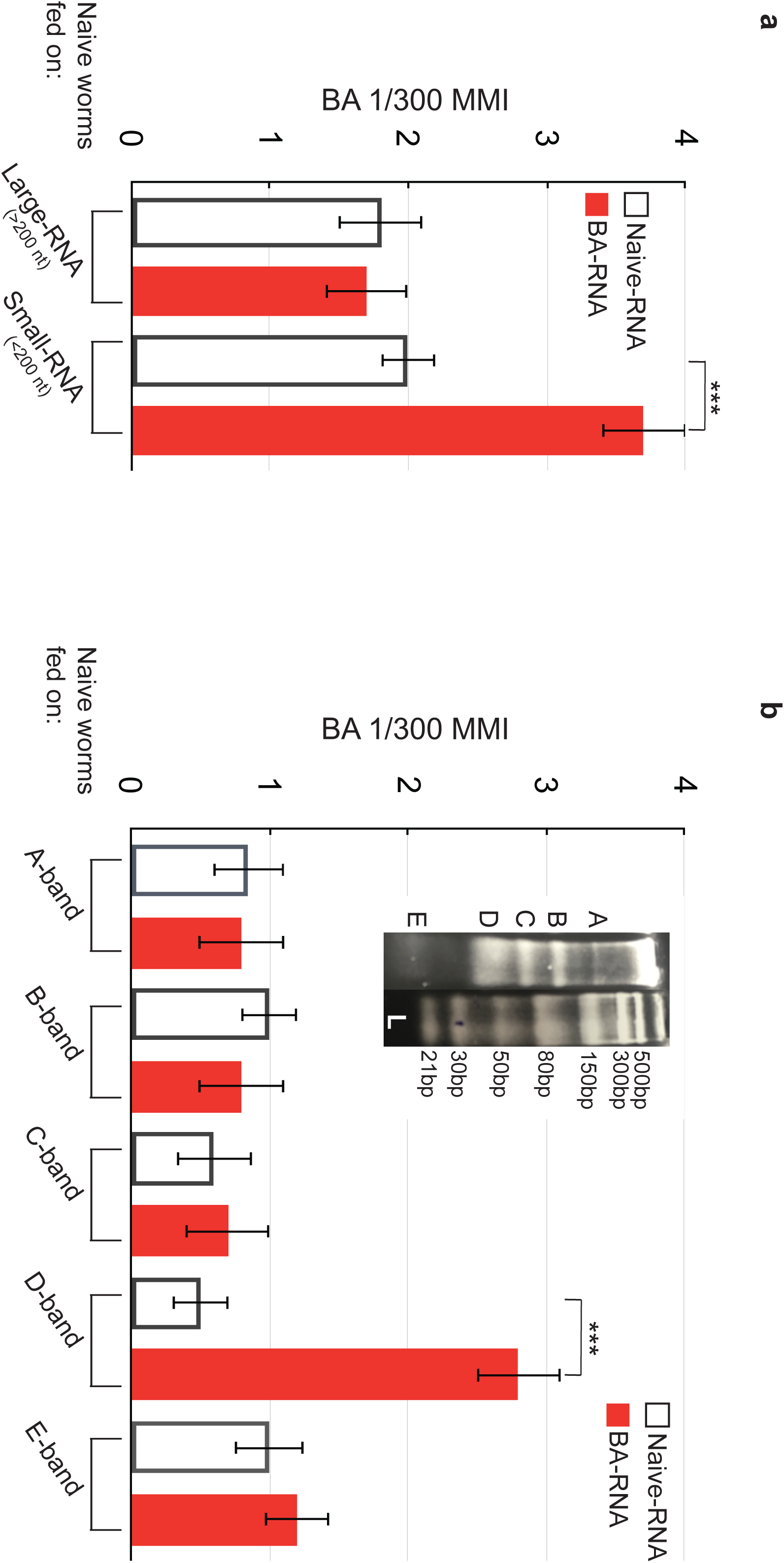
A specific population of small RNA transfers olfactory imprinting. **(Supp 1a)** RNA from BA-exposed and unexposed naive was fractionated into two populations: the larger than 200 nucleotides fraction (Large-RNA) and the smaller than 200 nucleotides fraction (Small-RNA). Worms fed on 10 ng of Large BA-RNA migrate in a BA gradient as worms fed on 10 ng of Large Naive-RNA. By contrast, worms fed on 10 ng of Small BA-RNA display enhanced BA responses, compared to worms fed on 10 ng of Small Naive-RNA or to worms fed on 10 ng of Large BA-RNA (experimental repeats > 4, ***p-value < 0.001). **(Supp 1b)** 3.5% low-melting agarose gels fractionate BA-RNA and Naive-RNA into five A to E discrete bands (left side of insert). Naive worms were fed on each of the five gel-purified NA-RNA and BA-RNA populations. Compared BA 1/300 MMI shows that only the BA-RNA band D is able to transfer a BA-imprint to naive (experimental repeats > 4, ***p-value < 0.001). (L), NEB double stranded RNA ladder.

**Supplemental Figure 2:**
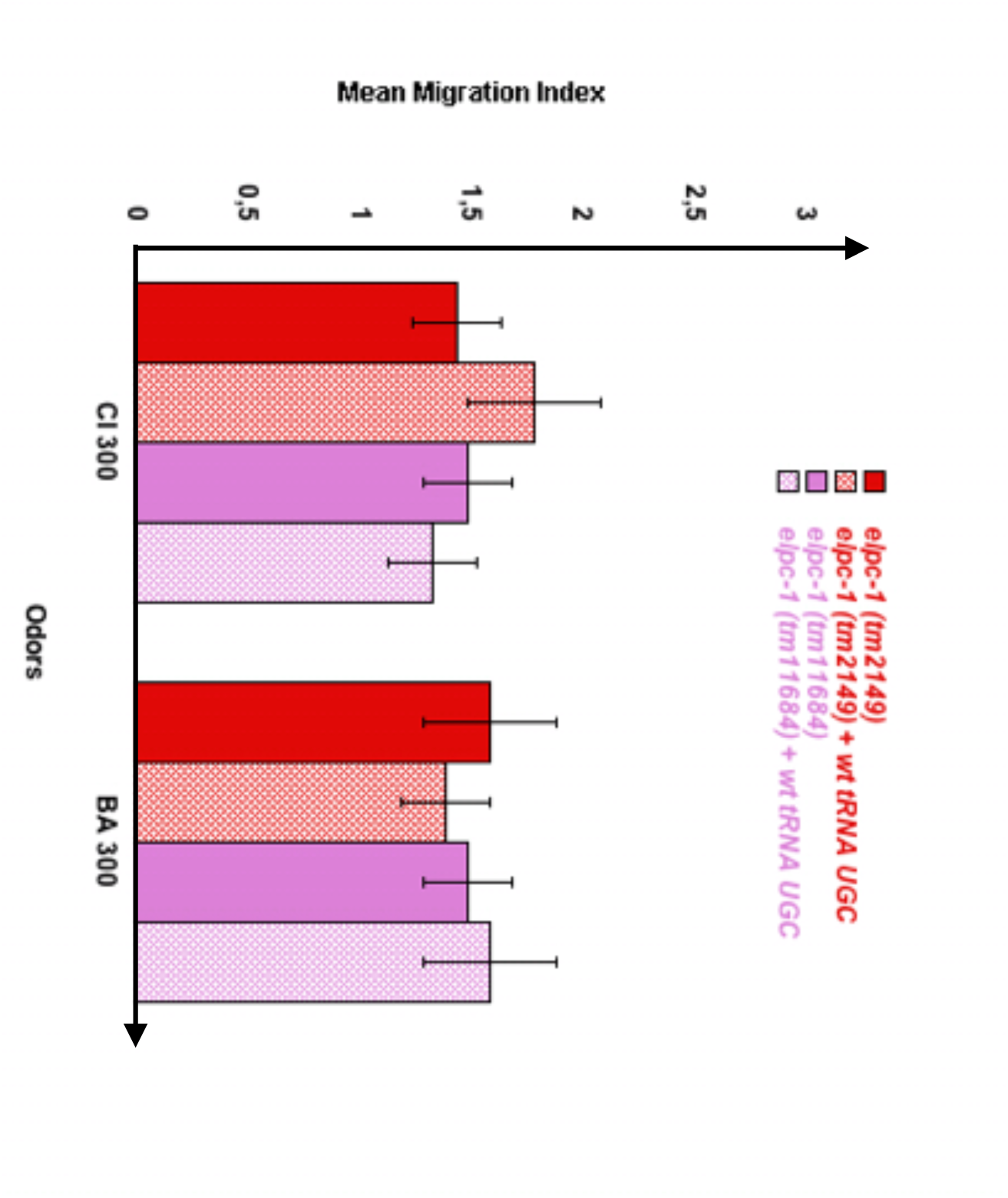
Wild-type tRNA^Ala^UGC has no effect on *elpc-1* mutant worms chemo-attraction. Compared chemo-attractive responses of naive or of wild-type tRNA^Ala^UGC fed *elpc-1 (tm2149)* and *elpc-1 (tm11684)* mutant worms. Mean Migration Indices (MMI) in gradients of Citronellol (CI), Benzaldehyde (BA) at the indicated dilutions, were determined as described (experimental repeats ≥ 4, **p-value < 0.01).

**Supplement Figure Method.**
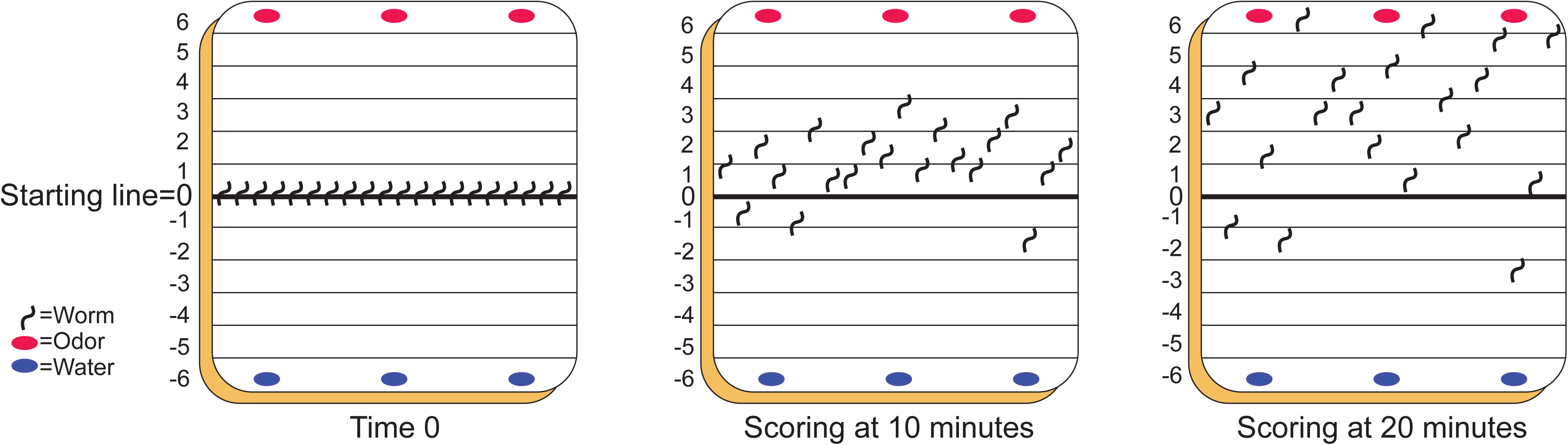

